# Co-Conservation of Synaptic Gene Expression and Circuitry in Collicular Neurons

**DOI:** 10.1101/2025.01.23.634521

**Authors:** Yuanming Liu, John A. McDaniel, Chen Chen, Lu Yang, Arda Kipcak, Elise L. Savier, Alev Erisir, Jianhua Cang, John N. Campbell

**Author notes:** equal-contributing senior authors (Jianhua Cang, John N. Campbell).

## Abstract

The superior colliculus (SC), a midbrain sensorimotor hub, is anatomically and functionally similar across vertebrates, but how its cell types have evolved is unclear. Using single-nucleus transcriptomics, we compared the SC’s molecular and cellular organization in mice, tree shrews, and humans. Despite over 96 million years of evolutionary divergence, we identified ∼30 consensus neuronal subtypes, including *Cbln2*+ neurons that form the SC-pulvinar circuit in mice and tree shrews. Synapse-related genes were among the most conserved, unlike neocortex, suggesting co-conservation of synaptic genes and circuitry. In contrast, cilia-related genes diverged significantly across species, highlighting the potential importance of the neuronal primary cilium in SC evolution. Additionally, we identified a novel inhibitory SC neuron in tree shrews and humans but not mice. Our findings reveal that the SC has evolved by conserving neuron subtypes, synaptic genes, and circuitry, while diversifying ciliary gene expression and an inhibitory neuron subtype.

## INTRODUCTION

Neurons are the most diverse cell type of the body, with thousands of molecular subtypes identified in the human brain so far (Siletti et al., 2023). Yet, much remains unknown about the evolutionary mechanisms shaping their forms and functions, including whether these differ across brain regions. Early brain research relied on comparative anatomy to explore evolutionary changes in brain size, organization, and cellular composition (Ramón y Cajal, 1995; Reiner, 1990). Later, electrophysiological techniques provided insights into neuronal function and connectivity in different species (Adrian, 1926; Roberts et al., 2022). As evolution works through changes in gene sequence and expression (Britten and Davidson, 1969; King and Wilson, 1975), many studies have applied DNA/RNA sequencing and other genetic and genomics tools and revealed the molecular mechanisms that differentiate brain development, size, organization, and function across species (Genome, 2009; Somel et al., 2013; Sun and Hevner, 2014; Wolfe and Li, 2003; Zhuang et al., 2024). More recently, single-cell transcriptomics has revolutionized the field by providing a high-throughput and real-time view of gene expression, cell type diversity, functional specialization, and developmental trajectories at cellular resolution (Caglayan et al., 2023; Hahn et al., 2023; Hain et al., 2022; Kebschull et al., 2020; La Manno et al., 2016; Network, 2021; Saunders et al., 2018; Shi et al., 2021).

Comparative single-cell transcriptomics has focused mostly on the neocortex, presumably because the most striking difference between the human brain and that of non-primates is the vast expansion of neocortical neurons and increased corticocortical connectivity (Lancaster, 2024). Several studies have shown that synapse-related genes have diverged across human, non-human primate, and rodent neocortex, consistent with differences in cortical circuitry (Berg et al., 2021; Hodge et al., 2019; Ma et al., 2022; Network, 2021; Peng et al., 2024; Sousa et al., 2017; Suresh et al., 2023). Thus, changes in synaptic gene expression during evolution may have facilitated synaptic rewiring of brain regions. However, it is not clear whether this principle also applies to other brain areas such as subcortical structures that evolved at a different rate compared to the cortex (Zhuang et al., 2024). Furthermore, it remains largely unknown how individual cell types and subcortical brain regions have diverged over evolution and whether synaptic genes have been conserved with synaptic circuitry.

To answer these fundamental questions, we studied a midbrain structure, the superior colliculus (SC). The SC is a major sensorimotor center that integrates multimodal sensory inputs to drive reflexive behaviors, as well as higher cognitive functions such as visual spatial attention and decision-making (Basso et al., 2021; Basso and May, 2017; Cang et al., 2024; Isa et al., 2021). In contrast to the neocortex, which differs dramatically across vertebrate species, the SC is considered as a conserved structure due to similarities across species in its laminar and topographic organization, cell morphology, and similar input and output connections (Basso et al., 2021; Liu et al., 2022). In this study, we specifically focused on the superficial visual layers of the SC (sSC). We used single-nucleus transcriptomics, RNA fluorescence in situ hybridization, and retrograde tracings to compare the sSC’s molecular and anatomical organization between mice (*M. musculus*) and northern tree shrews (*T. belangeri*), species which diverged in evolution over 96 million years ago (Fan et al., 2013). The tree shrew is a close relative of primates and an emerging model in neuroscience research (Campbell, 1966; Savier et al., 2021; Yao et al., 2024). To determine whether the similarities and differences we found between mouse and tree shrew SC also extended to human SC, we incorporated human single-nucleus transcriptomics data into our comparative analysis.

Our results identify ∼30 consensus molecular subtypes of SC neurons across the three species. Notably, in contrast to previous comparative studies of the cortex, genes for synapse organization and function are among the most similarly expressed across mouse, tree shrew, and human SC neurons. For example, a specific subtype of glutamatergic neuron, marked by synaptic molecule *Cbln2+* in all 3 species, provide most of the SC’s projection to the pulvinar in both mice and tree shrews, indicating a co-conservation of synaptic gene expression and synaptic circuitry. Interestingly, transcripts related to cilia function are among the most divergent across the mouse, tree shrew, and human SC, suggesting that the primary cilium, a signaling hub for neurons, might have been a hot spot during SC evolution. Finally, we detected an inhibitory neuron subtype in the tree shrew and human SC but not in the mouse SC, suggesting divergence of inhibitory neuron subtypes during evolution. Together, our results demonstrate that the SC evolved through distinct mechanisms compared to the neocortex. Our results also shed light on how SC cell types have diversified, revealing inhibitory neuron subtypes which diverged highly during evolution and an enrichment of transcripts related to primary cilium function in higher order species.

## RESULTS

### Cellular composition and anatomy are conserved between mouse and tree shrew SC

We profiled the genome-wide mRNA expression of female and male tree shrew superficial superior colliculus (sSC) using single-nucleus RNA sequencing (snRNA-seq) and integrated the dataset with our recently published one from mouse sSC (Liu et al., 2023) (Figure S1A; hereafter referred to as ‘TM dataset’). We clustered 69,492 cells into 31 consensus cell types based on expression of high variance genes (Figure 1A, S1B). Neurons comprised the largest proportion and highest diversity of sSC cells in both species (Figure 1B, S1C). Compared with the mouse sSC, the tree shrew sSC contained a slightly lower proportion of neurons but a higher proportion of oligodendrocyte lineage cells (i.e., oligodendrocytes and oligodendrocyte precursor cells).

**Figure 1.**
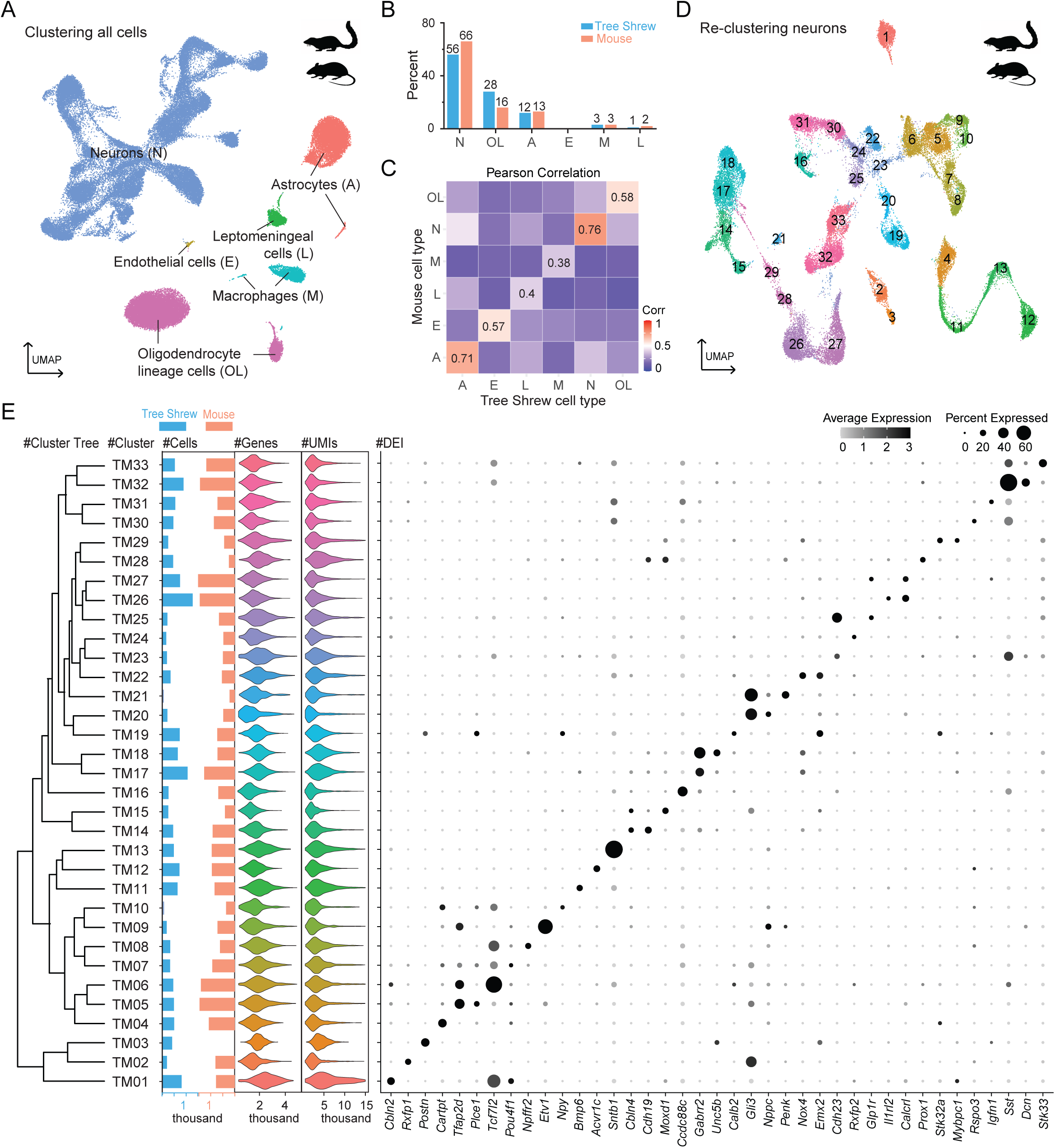
Transcriptomically defined cell types and neuronal subtypes in the mouse and tree shrew superficial superior colliculus (sSC) A. UMAP (uniform manifold approximation and projection) plot of integrated cells from the mouse and tree shrew sSC, clustered by expression of highly variable genes and colored by cell type. Each dot represents a cell. B. Percentage of cells for each cell type in the two species. C. Heatmap showing Pearson correlation coefficient of gene expression for each cell type between mice and tree shrews. D. UMAP plot of integrated neurons after filtering, colored by neuronal subtype identity. E. Dendrogram illustrating the transcriptomic relatedness of 33 neuronal subtypes, followed by cluster names, cell counts per subtype in tree shrews and mice, the number of detected genes and transcripts per subtype, and a plot showing average expression and percentage of cells expressing the marker genes selectively enriched for each subtype.

We then compared the transcriptomes of each cell type across sex and species. Across all cell types, neurons showed the highest correlation between tree shrew and mice, followed by astrocytes (Figure 1C, S1D). In contrast, macrophages displayed the greatest divergence in gene expression (Figure 1C, S1D). Additionally, when comparing gene expression between sexes within species, neurons were the most conserved cell type between females and males (Figure S1E-H). Leptomeningeal cells and endothelial cells were the most sexually dimorphic in mice and tree shrews, respectively (Figure S1E-H). Together, our results indicate that neurons are the most conserved across species and between sexes.

To compare neuronal subtypes between species, out of the 69,492 total cells, we subsetted and re-clustered 42,940 neurons into 33 distinct clusters and identified cluster-enriched genes (Figure 1D-E). This clustering of mouse and tree shrew neurons in the sSC was driven by their molecular similarity and present across a range of clustering parameters (Figure S2A-H). We next assessed conservation of the 33 neuronal subtypes between the two species (Figure 2A-B). Both species had more clusters of inhibitory neurons than excitatory neurons, based on their relative expression of neurotransmitter phenotype genes *Gad2* (glutamate decarboxylase 2) and vGluT2 (vesicular glutamate transporter 2, gene *Slc17a6*; Figure 2B), respectively. Both species also displayed a similarly high proportion of inhibitory neurons, which constituted an average of 64% of neurons in the tree shrew and mouse integrated dataset, which were validated by RNA fluorescence in situ hybridization (RNA FISH; Figure 2C-E). Inhibitory neurons were enriched in the most superficial layers of the SC, whereas excitatory neurons were more prevalent in deeper layers (Figure 2F). Notably, the overall density of neurons was roughly twice as high in mice than in tree shrews, with the difference largely uniform along SC depth (Figure 2G). These results suggest that sSC neurons have similar composition and spatial organization in the two species but different density and varying proportions of individual transcriptomic clusters.

**Figure 2.**
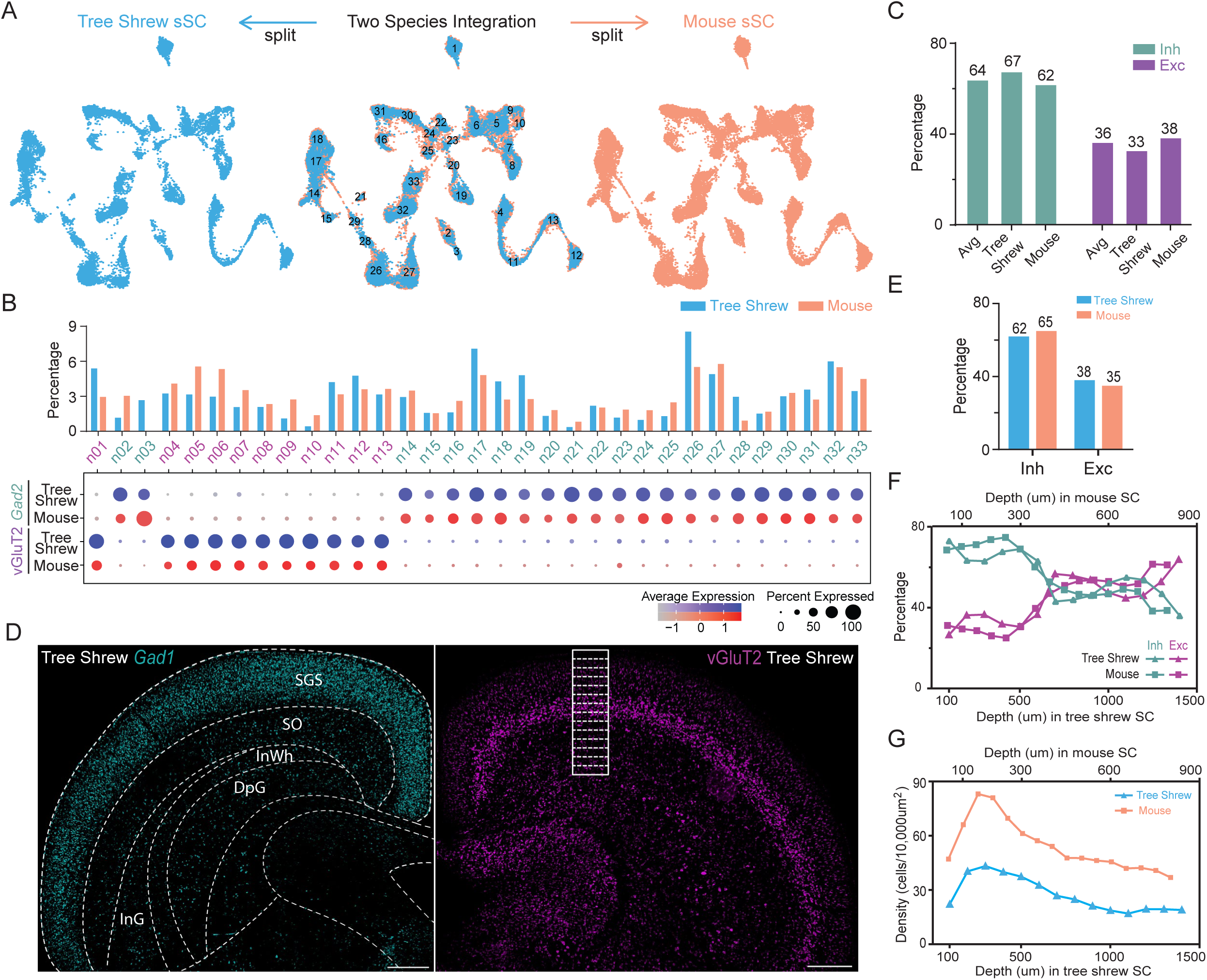
Neuronal composition and spatial organization in the mouse and tree shrew sSC. A. Visualization of neurons by species on the integrated UMAP (middle), with tree shrew neurons (left) and mouse neurons (right) shown in the same UMAP space. B. Percentage of neurons in each neuronal subtype by each species (top), along with their expression of inhibitory (*Gad2*) and excitatory (vGluT2, bottom) markers. C. Percentage of inhibitory and excitatory neurons in the integrated dataset. D. Fluorescence *in situ* hybridization (FISH) of *Gad1* and vGluT2 expression in the tree shrew SC. Left, laminar organization of SC is indicated according to the stereotaxic coordinates of the tree shrew brain (left). Right, a 1400 μm × 400 μm white rectangle is used to quantify expression and is further divided into 14 subregions along the SC depth. Scale bars: 500 μm. E. Percentage of inhibitory and excitatory neurons labeled by FISH within the tree shrew and mouse sSC. n = 3 animals each. F. Percentage of inhibitory and excitatory neurons along the SC depth in tree shrews (triangles) and mice (squares). G. Neuronal density along the SC depth in tree shrews and mice.

### Synapse-related genes are conserved between mouse and tree shrew sSC neurons

Given the close correspondence of sSC cell types and neuronal subtypes we observed between mice and tree shrews, we investigated the conservation of gene expression between the two species. For each orthologous gene, we calculated expression differences between mouse and tree shrew sSC in log2 fold-change (hereafter referred to as ‘species specificity index (SSI)’; Figure 3A). The SSI values were normally distributed around 0, indicating that most genes are expressed at similar levels between species (Figure S3A). We then performed Gene Ontology (GO) analysis on the 1,600 genes with the smallest absolute SSI values to categorize the genes with the most conserved expression levels. These genes were enriched with GO terms associated with synaptic functions and nervous system development, including: postsynapse organization; synapse assembly; regulation of transporter activity; dendrite development; and the regulation of neurogenesis and gliogenesis (Figure 3B). Synapse-, postsynapse-, and projection organization-related genes, for instance, had a higher percentage of SSI values close to 0 than general genes, indicating greater conservation (Figure 3C, S3B). Among postsynapse-related genes, 52% had absolute SSI values lower than 1, compared to 45% for synapse-related genes and 34% for all genes, suggesting that the expression of postsynapse-related genes is more conserved than other synaptic genes (Figure S3C). Repeating our analysis with a synapse-focused GO method, SynGO (Koopmans et al., 2019), confirmed enrichment of genes for synaptic processes and organization among the most conserved between mouse and tree shrew sSC (Figure 3D, S3D).

**Figure 3.**
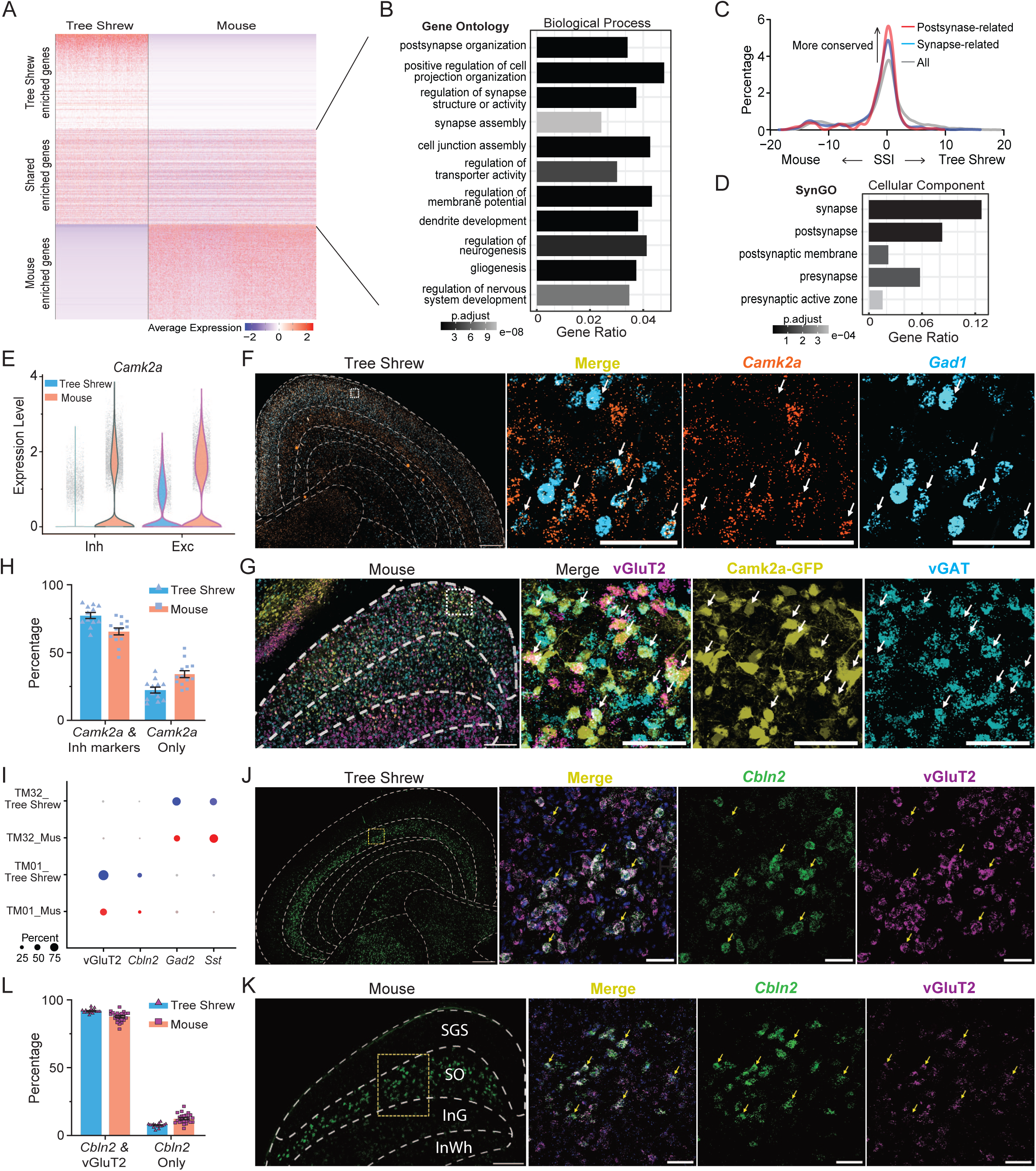
Synapse-related genes are conserved between mouse and tree shrew sSC neurons. A. Heatmap showing the expression of the top 500 genes enriched in tree shrews, mice or shared by the two species. B. Enriched Gene Ontology (GO) terms related to biological processes for the top 1600 shared genes. C. Percentage distribution of species specificity index (SSI) for all genes, synapse-related genes, and postsynapse-related genes. Positive x-axis values: higher gene expression in tree shrews; negative: higher in mice. D. SynGO enrichment analysis for the top 1600 shared genes in the aspect of cellular components. E. Violin plot showing *Camk2a* expression in inhibitory and excitatory neurons by species in the snRNA-seq dataset. Each dot represents an individual neuron. F. FISH of *Camk2a* and *Gad1* transcripts in the tree shrew SC. The area in the white square is shown at a higher magnification on the right. White arrows point to example cells showing co-localization. Scale bars: 500 μm (left) and 50 μm (right). G. FISH of vGAT and vGluT2 expression in the mouse SC, with neurons labeled by AAV-Camk2a-GFP. White arrows indicate example cells where GFP+ neurons co-localized with vGAT or vGluT2. Scale bars: 200 μm (left) and 50 μm (right). H. Percentage of *Camk2a+* cells co-localized with inhibitory markers in mice and tree shrews based on RNA FISH. Mean ± SEM. n = 12 ROIs in tree shrews and 13 ROIs in mice; n = 3 animals each. I. Example clusters showing co-expression of cluster markers related to synaptic functions and inhibitory or excitatory markers. J -K. FISH of *Cbln2* and vGluT2 expression in tree shrews (J) and mice (K). Scale bar, 500 μm (J, left), 200 μm (K, left), and 50 μm (right). L. Percentage of *Cbln2*+ cells co-expressing vGluT2. Mean ± SEM. n = 16 ROIs in tree shrews and 24 ROIs in mice; n = 3 animals each.

We then examined the anatomical expression of conserved synapse-related genes in mouse and tree shrew sSC. For example, expression of calcium–calmodulin dependent protein kinase II alpha (*Camk2a*), which is crucial for synaptic plasticity (Yasuda et al., 2022), was detected in both inhibitory and excitatory neurons across most subtypes in both species (Figure 3E, S3E). This contrasts with the *Camk2a* expression in the hippocampus and cortex where it is abundant in glutamatergic neurons. To confirm this, we co-localized *Camk2a* with the inhibitory neuron marker *Gad1* in the tree shrew SC and cortex using RNA FISH. The majority of *Camk2a+* cells at the superficial SC in fact co-expressed *Gad1* (77.48 ± 2.2%), with this co-expression decreasing with depth in the SC (Figure 3F, H, S3F). In contrast, very few *Camk2a*+ cells in the tree shrew cortex co-expressed *Gad1* (Figure S3G, I). In the mouse SC, *Camk2a* expression was widespread and strong, making it difficult to distinguish its expression in neurons. We thus injected AAV-Camk2a-GFP virus into the mouse SC and cortex and co-localized GFP with the inhibitory neuron marker vGAT or the excitatory neuron marker vGluT2 by RNA FISH (Figure 3G). More than half of GFP+ cells in the mouse SC were labeled with vGAT transcripts, and a similar pattern was observed for vGluT2 expression (Figure 3G-H). This was again different from the cortex, where nearly all GFP+ cells were negative for *Gad1* expression (Figure S3H-I). These results confirm the snRNA-seq analysis result that both inhibitory and excitatory neurons in the sSC express *Camk2a* in both species.

Additionally, one synapse-related gene, cerebellin 2 (*Cbln2*), which encodes a secreted neuronal glycoprotein essential for synapse formation and maintenance (Dai et al., 2022; Shibata et al., 2021; Sudhof, 2023), showed enriched expression in an excitatory cluster of the integrated atlas, TM01 (Figure 3I). RNA FISH validated co-expression of *Cbln2* and vGluT2 in both mouse and tree shrew sSC (Figure 3J-L). Furthermore, *Cbln2* expression was restricted to the lower SGS/SO sublayer of both species, indicating that *Cbln2*+ cells are similarly located in the tree shrew and mouse sSC (Figure 3J-K). Another synapse-related gene, somatostatin (*Sst*), which encodes a neuropeptide involved in synaptic communication, had enriched expression in an inhibitory neuron subtype, TM32, in both the mouse and tree shrew sSC (Figure 3I). RNA FISH validated co-expression of *Sst* and *Gad1* in the tree shrew SC and cortex, where almost all *Sst*+ cells contained *Gad1* transcripts, consistent with our previous observations in mice (Liu et al., 2023) (Figure S3J-L). Furthermore, *Sst*+ cells were similarly distributed in both species, localized in the lSGS and SO sublayers (Figure S3J). In summary, these results demonstrate that synapse-related genes, the neurotransmitter of each subtype, and the spatial organization of at least some neuronal subtypes are conserved between the tree shrew and mouse sSC.

### A conserved subtype expressing Cbln2 projects to the pulvinar in both mice and tree shrews

Given the conserved molecular and anatomical features of the sSC between mice and tree shrews, we next compared the neural circuitry of one common neuronal subtype. In mice, *Cbln2* (a marker of TM01) is known to mark the so-called wide field vertical cells that project to the pulvinar (also known as the lateral posterior nucleus of the thalamus) (Choi et al., 2023; Xie et al., 2021). To confirm this, we first retrogradely labeled pulvinar-projecting neurons by injecting retrograde AAVrg-Cre into the pulvinar of H2B-TRAP mice, which Cre-dependently express a nuclear fluorescent protein (Figure 4A.1). This labeled a subset of SC neurons (Figure 4A.2). We isolated cell nuclei from the labeled neurons, transcriptionally profiled them, and clustered them into nine subtypes, including seven neuronal subtypes, accounting for 93.1% of all labeled cells, as well as two glial cell subtypes (Figure 4A.4, S4A-B).

**Figure 4.**
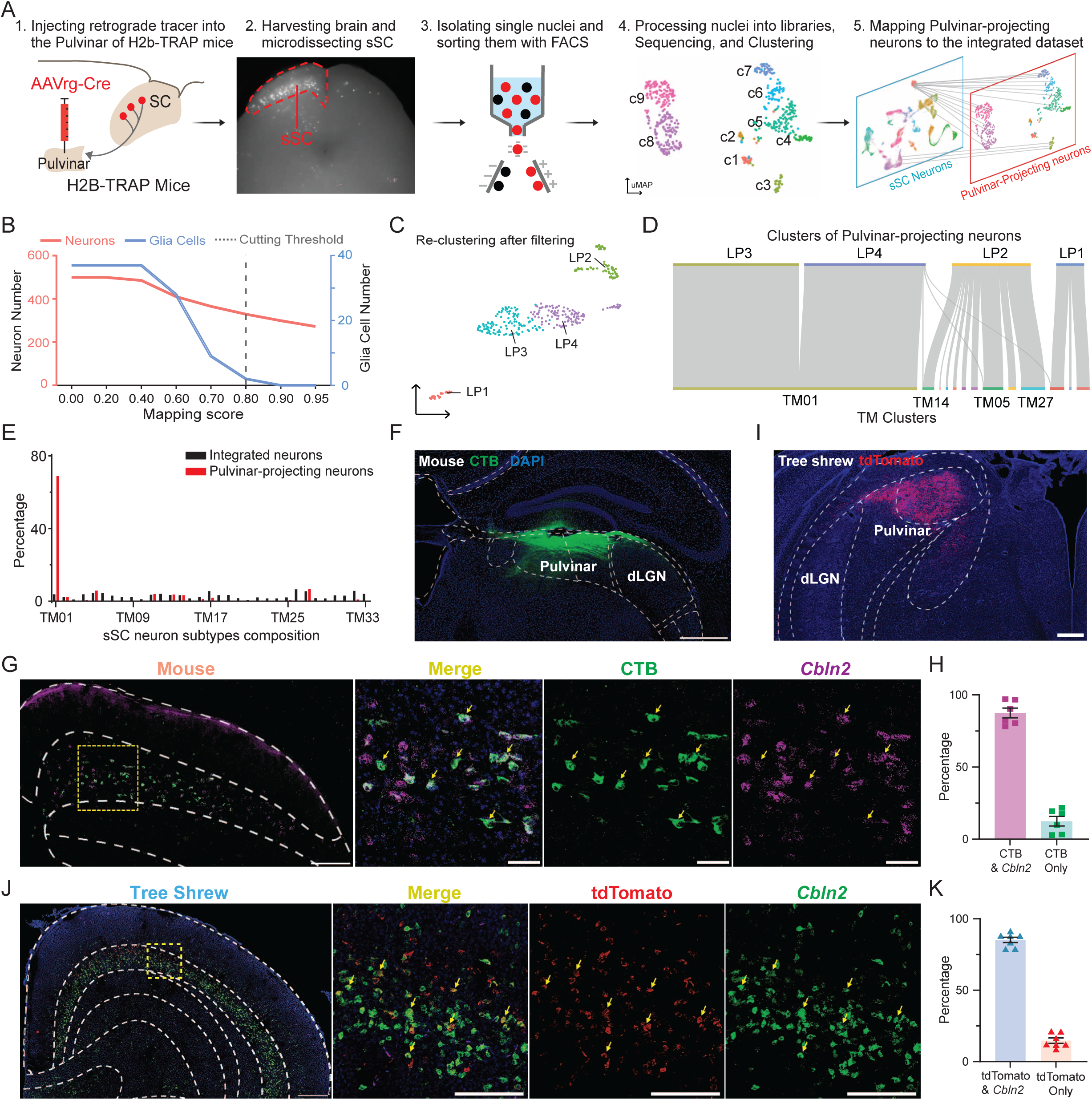
A conserved SC neuron subtype expressing Cbln2 projects to the pulvinar in both mice and tree shrews. A. Workflow for sequencing and annotating pulvinar-projecting neurons in the mouse SC. A retrograde tracing virus, AAVrg-Cre, was injected into the pulvinar of H2B-TRAP mice (A1) to retrogradely express Cre in the projecting neurons, thereby inducing H2B-mCherry expression (A2). The single nuclei of these projecting neurons were sorted based on mCherry expression (A3), sequenced, visualized in a UMAP plot (A4), and annotated by mapping to the integrated dataset (A5). n=9 mice. B. Number of neurons (red, left axis) and glia cells (blue, right axis) after filtering based on maximum mapping score during label transfer annotation. C. UMAP of pulvinar-projecting neurons that passed the filtering threshold in B. D. Sankey plot visualizing the mapping of pulvinar-projecting clusters to the integrated dataset. E. Percentage of neurons in each of the 33 integrated subtypes (black) compared to the percentage of LP-projecting neurons mapped to these subtypes (red). F. Expression of CTB488 tracer injected into the mouse LP, with the structure outline depicted according to the Allen Mouse Brain Atlas. Scale bar, 500 μm. G. FISH of *Cbln2* expression in the mouse SC with pulvinar-projecting neurons labeled with CTB488. White arrows indicate co-localization. Scale bars, 200 μm (left) and 50 μm (right). H. Percentage of CTB-labeled neurons expressing *Cbln2* transcript. Mean ± SEM. n = 6 images; n = 3 animals I. Expression of a retrograde tracer, AAVrg-tdTomato, injected into the tree shrew pulvinar. Example image shows injection in dorsal pulvinar. Scale bar, 500 μm. J. FISH of *Cbln2* and tdTomato transcripts in the tree shrew SC. Scale bars, 500 μm (left) and 100 μm (right). K. Percentage of tdTomato+ neurons co-expressing *Cbln2*. Mean ± SEM. n = 7 images; n = 3 animals.

To identify the retrogradely labeled SC cells, we mapped their transcriptomes to the TM dataset. We filtered out cells mapping with a confidence score less than 0.8 (out of 1.0), based on the confidence scores at which almost no glia map to the reference dataset, which contained only neurons (Figure 4A.5, B). A total of 66% of neurons (329 out of 499) met this criterion. About 70% of these pass-filter neurons mapped to TM01, a percentage 18-fold higher than would be expected based on random mapping (Figure 4C-E) and largely independent of the confidence filter (Figure S4C-D). These results thus indicate that the pulvinar-projecting sSC neurons in mice are primarily of the subtype TM01.

To validate these findings in vivo, we injected the retrograde tracer, Cholera Toxin Subunit B (CTB), into the mouse pulvinar and co-localized its labeling with *Cbln2* RNA FISH. CTB injection into the pulvinar labeled neurons in the SO sublayer of the SC (Figure 4F-G). Most of the labeled neurons (87.5 ± 3.4%) contained *Cbln2* transcripts, consistent with our snRNA-seq results (Figure 4H). To investigate whether the pulvinar in the tree shrew receives input from the same subtype of SC neurons, we injected a retrograde, tdTomato-expressing adeno-associated virus (AAV) into the tree shrew pulvinar and co-localized tdTomato with *Cbln2* transcripts in the SC by RNA FISH (Figure 4I-J). Robust tdTomato expression was detected in the lower SGS/SO sublayer of the tree shrew sSC, with most of these cells (85.3 ± 1.9%) co-expressing *Cbln2* (Figure 4K). Together, our results demonstrate that a conserved molecular neuron type, TM01, is the major source of synaptic output to the pulvinar in both tree shrew and mouse sSC.

### Primary cilium-related genes are enriched in the tree shrew sSC

Evolution likely works in part through changes in gene identities and expression levels (Britten and Davidson, 1969; King and Wilson, 1975). We therefore investigated the differences in gene expression between mouse and tree shrew SC neurons to identify molecular mechanisms of their evolution. First, we performed GO analysis on the top 1600 genes enriched in the mouse sSC based on SSI values. In terms of biological processes, these mouse-enriched genes were associated with chromosome organization, RNA processing, organelle localization, and cellular metabolic processes (Figure 5A). On the other hand, our GO analysis of the top 1600 tree shrew-enriched genes in the biological process category revealed enrichment of genes for cilium movement, sperm motility, second messenger-mediated signaling, adenylate cyclase (cAMP)-modulating G protein-coupled receptor (GPCR) signaling pathways, calcium-mediated signaling, and lipid transport (Figure 5B). We also examined enriched GO terms in the category of cellular components and found that they included motile cilia, secretory granules, collagen-containing extracellular matrix, and acrosomal vesicles (Figure 5C). Reciprocally, genes related to cilium movement, secretory granule, and regulation of cilium movement showed a lower percentage of SSI values near 0 but a higher percentage of positive SSI values than general genes, indicating enrichment in the tree shrew sSC relative to mice (Figure 5D, S5A).

**Figure 5.**
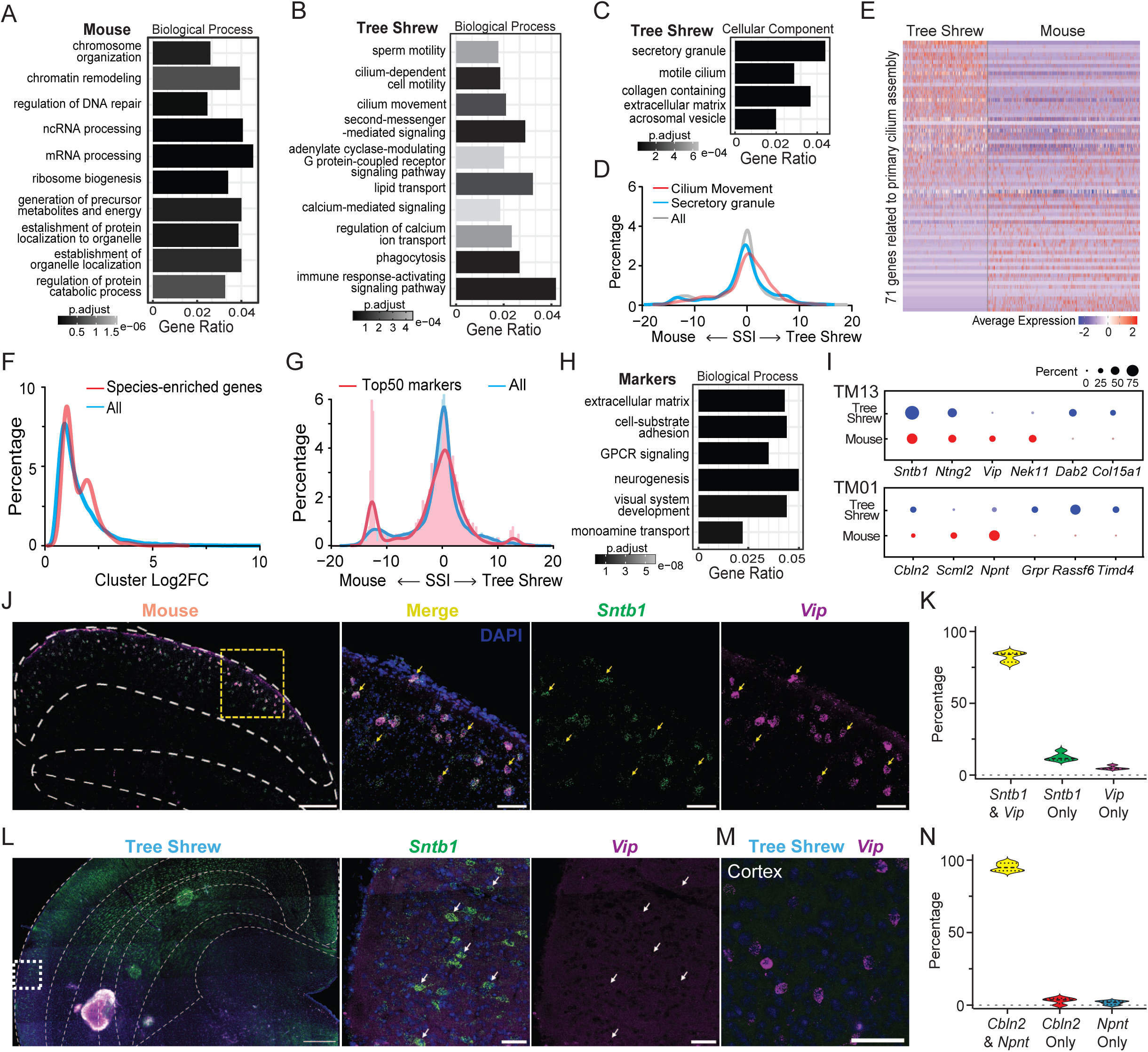
Primary cilium-related genes and cluster markers are divergent between mouse and tree shrew sSC neurons. A. GO enrichment analysis of the top 1600 mouse-enriched genes. B. GO enrichment analysis of the top 1600 tree shrew-enriched genes. C. GO enrichment analysis for cellular components among the top 1600 tree shrew-enriched genes. D. Percentage distribution of SSI of all genes, cilium movement-related genes, and secretory granule-related genes. E. Heatmap illustrating the expression of 71 genes related to primary cilium assembly across neurons in the dataset. F. Percentage distribution of the maximum log2 fold change of gene expression between clusters (cluster Log2FC) for all genes and the top 1000 species-enriched genes, with 500 from each species. G. Percentage distribution of SSI for all genes and the cluster markers (top 50 genes for each cluster). H. GO enrichment analysis on the cluster markers (top 50 genes for each cluster). I. Example clusters showing markers either shared by the two species or enriched in one species. J. FISH of *Sntb1* and *Vip* in the mouse SC. Scale bars, 200 μm (left) and 20 μm (right). K. Percentage of mouse sSC neurons expressing *Sntb1* and *Vip*, *Sntb1* only, or *Vip* only. Mean ± SEM. n = 4 images; n = 2 animals. L. FISH of *Sntb1* and *Vip* in the tree shrew SC. Scale bars, 500 μm (left), and 20 μm (right). M. FISH of *Vip* in the tree shrew cortex. Scale bars, 20 μm (right). N. Percentage of mouse sSC neurons expressing *Cbln2* and *Npnt*, *Cbln2* only, or *Npnt* only. Mean ± SEM. n = 5 images; n = 3 animals.

Although genes related to motile cilia and their movement were enriched in tree shrew sSC neurons, motile cilia have not been reported in SC neurons. Instead, primary cilia, which are antenna-like sensory organelles, are prevalent in neurons. Primary cilia are enriched with receptors, particularly GPCRs, to receive extracellular signals, and they use signaling cascades to amplify and transduce signals to the soma via secondary messengers such as cAMP and calcium (Jurisch-Yaksi et al., 2024; Wu et al., 2024; Youn and Han, 2018). Therefore, the motile cilia-related genes identified by our GO analysis are likely related to the function of the neuronal primary cilia. Concurrently, approximately one-third of genes related to primary cilium and primary cilium assembly were enriched in tree shrew with SSI values greater than 1 (Figure 5E, S5B-C). Our results therefore suggest that tree shrew and mouse sSC diverged highly in terms of their expression of primary cilia-related genes.

### Cluster marker genes are divergent between mice and tree shrews

We next explored whether the species-enriched genes were more likely to occur within certain neuronal subtypes. We calculated the expression differences of each orthologous gene between clusters in log2 fold-change (Cluster Log2FC). Compared to all genes, the species-enriched genes showed a higher percentage of larger Cluster Log2FC values, suggesting a cluster specificity among some divergent genes (Figure 5F). On the other hand, the top 50 markers for each cluster together displayed a lower percentage of SSI values near zero and a higher percentage of larger absolute SSI values than all genes, providing complementary evidence that certain cluster markers tend to differ between species (Figure 5G). Furthermore, GO analysis of these top 50 markers revealed functions associated with cell-substrate adhesion, neurogenesis, neurotransmission, and GPCR signaling pathways (Figure 5H). Together, our findings suggest that even though the neuron types are remarkably conserved in mouse and tree shrew sSC, many marker genes of these molecular neuron types are different in the two species.

To validate these observations, we identified markers for some neuron clusters and confirmed their convergent and divergent expression in the mouse and tree shrew sSC with RNA FISH. For instance, syntrophin beta 1 (*Sntb1*), which encodes a cytoskeletal protein, showed enriched expression in an excitatory cluster, TM13, in both species (Figure 5I, S5D). For the same cluster, vasoactive intestinal peptide (*Vip*), which encodes a signaling neuropeptide, was a marker gene in the mouse, but not expressed in tree shrew sSC (Figure 5I). RNA FISH validated the co-localization of *Sntb1* and *Vip* in the mouse sSC, but detected only *Sntb1*, not *Vip* in the tree shrew sSC (Figure 5J-L). In contrast, *Vip* transcripts were detected in the cortex of the same tree shrews, confirming that the lack of *Vip* expression in the tree shrew sSC was not due to technical issues (Figure 5M). Another sSC neuronal subtype, TM01, contained *Cbln2* transcripts in both species but expressed nephronectin (*Npnt*), which encodes an extracellular binding protein (Tsai et al., 2022), in mice but not in tree shrews (Figure 5N, S5E-F). Furthermore, the spatial organization of cluster TM13 differed between species, in that *Sntb1*+ cells were restricted to the surface of the mouse SGS, whereas in the tree shrew, they were primarily located in the middle of the SGS (Figure 5J, L). Overall, our results highlight that cluster markers are more divergent during evolution.

### An inhibitory neuron subtype is present in the tree shrew but not the mouse sSC

Our clustering analysis identified one subtype of inhibitory neurons, TM03, present in the sSC of tree shrews but not mice (Figure 2B, 6A). This previously uncharacterized subtype was closely related to another inhibitory subtype, TM02, which is present in both species (Figure 1C). The presence of the TM03 neuron cluster and its close relatedness to TM02 were stable across clustering parameters, numbers of cells, and dataset normalization and integration methods, indicating that it is not an artifact of data processing (Figure 6B-C, S6A-I). To further determine the genes driving divergence between the two clusters, we performed differential gene expression analysis between TM03 and TM02 (Figure 6D). We then performed GO analysis on these two sets of enriched genes separately. Notably, genes enriched in both TM03 and TM02 are associated with functions such as cell-cell adhesion, synapse assembly, cell junction assembly, cell projection, and dendrite development, suggesting a distinct set of synapse-related genes drive the emergence of TM03 in the tree shrew SC (Figure 6E-F). For instance, TM02 and TM03 differed in their expression of members of the cadherin family, cell-cell adhesion proteins which specify synaptic connectivity (de Wit and Ghosh, 2016): expression of *Cdh2*, *Cdh6*, and *Cdh9* was more enriched in TM03, whereas *Cdh12* and *Cdh18* showed higher expression in TM02, raising the possibility that TM02 and TM03 connect with different synaptic partners. Other transcriptional differences imply that TM03 and TM02 may respond differently to fast acting neurotransmitters. For instance, relative to TM03, TM02 showed higher expression of the GABA receptor subunits *Gabrb2*, *Gabra1*, and *Gabrg2*, the AMPA receptor subunit gene *Gria1*, and the ionotropic glutamate receptor gene *Grid2*. In summary, our observations suggest that TM02 and TM03 are related neuronal subtypes that may have diverged from a common cellular ancestor during sSC evolution.

**Figure 6.**
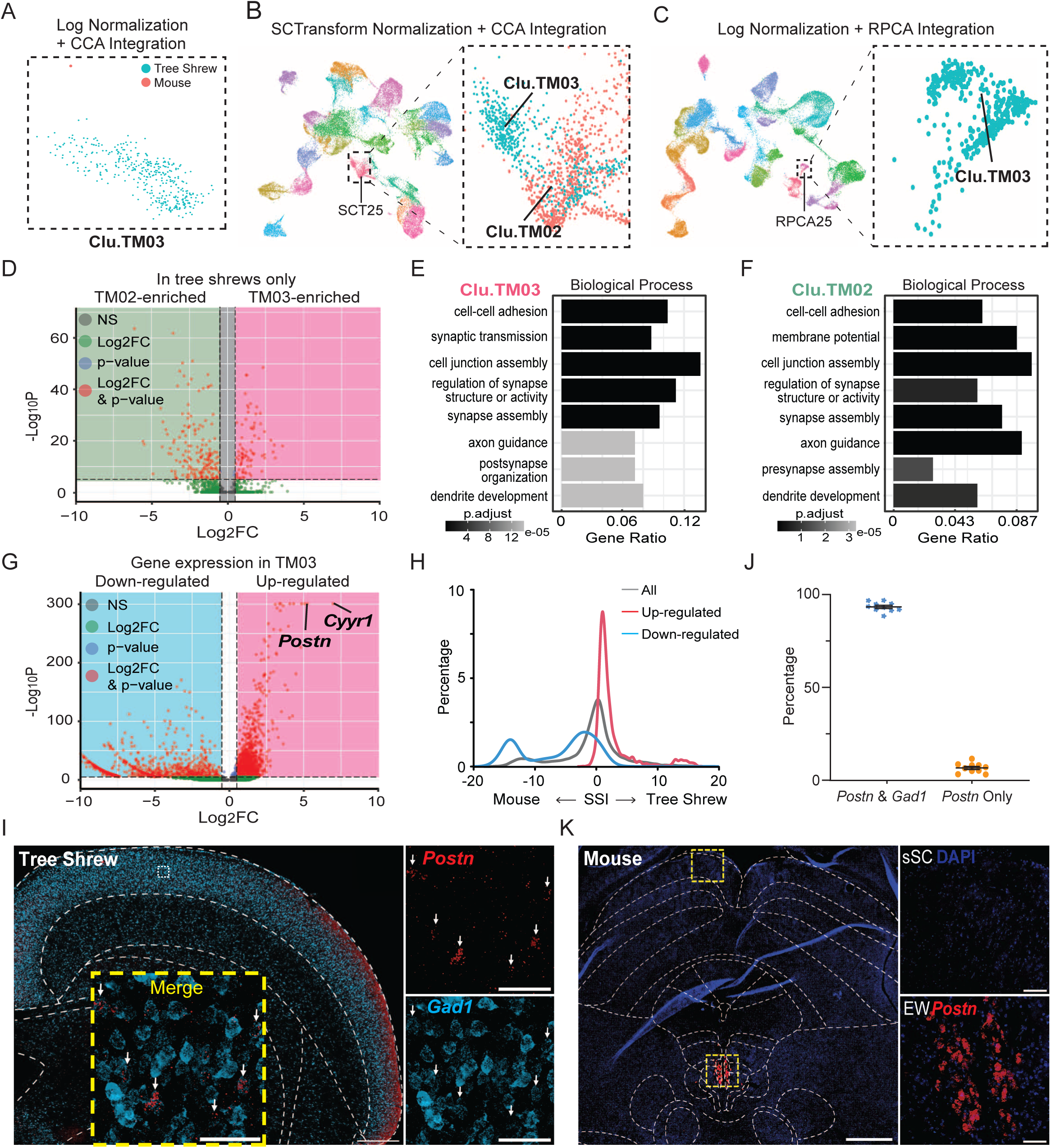
An inhibitory neuron subtype is present in the tree shrew sSC but not the mouse sSC. A. Visualization of the tree shrew-enriched subtype, Clu.TM03, by species on the integrated UMAP space using log normalization and anchor-based CCA integration. B. UMAP plot of integrated dataset using SCTransform normalization and CCA integration. Cluster SCT25 is shown at higher magnification on the right. Left, neurons are colored by cluster identity. Right, neurons are colored by species with a subpopulation in the top left identified as Clu.TM03. C. UMAP plot of integrated dataset using log normalization and RPCA integration. Cluster RPCA25 is shown at higher magnification on the right. Right, neurons are colored by species with all identified as Clu.TM03. D. Volcano plot showing genes with enriched expression in TM03, TM02, or shared by the two clusters. Genes are colored based on thresholds: |log₂FC| > 0.5 and p-value < 1 × 10⁻⁵. Red indicates genes meeting both criteria, blue for log2FC only, green for p-value only, and grey for neither. E. GO enrichment analysis of the genes enriched in Clu.TM03. F. GO enrichment analysis of the genes enriched in Clu.TM02. G. Volcano plot illustrating genes with enriched expression within or outside TM03. The top two genes, identified by the highest log2 fold change (log2FC) and smallest p-values, are labeled with their respective symbols. H. Percentage distribution of SSI for all genes, genes enriched within TM03, and genes enriched in other clusters. I. FISH of *Postn* and *Gad1* in the tree shrew SC. The area in the yellow square is shown at a higher magnification at the bottom, with split channels displayed on the right. Scale bars, 500 μm (left) and 50 μm (bottom and right). J. Percentage of *Postn*+ neurons co-expressing *Gad1* in the tree shrew SC. Mean ± SEM. n = 10 ROIs; n = 2 animals. K. FISH of *Postn* in a mouse coronal section. The area in the top yellow square is magnified at top right, whereas the bottom square is shown at bottom right. Scale bars, 500 μm (left) and 50 μm (right).

We identified genes with enriched expression in TM03 relative to all other clusters and performed GO analysis to explore their potential functions (Figure 6G). The top GO terms included GTPase mediated signal transduction, synaptic organization, and dendrite development (Figure S6J). Moreover, transcripts enriched in this neuron cluster relative to the others tended to be highly specific to tree shrews, whereas transcripts enriched in other clusters tended to show greater expression in the mouse sSC (Figure 6G-H). These results suggest that a species-specific regulatory mechanism of gene expression, involving both upregulation and downregulation, may contribute to the emergence of this distinct inhibitory subtype.

To anatomically map the TM03 neuron cluster in vivo, we selected a top marker gene, *Postn*, and performed RNA FISH (Figure 6G, S6K). *Postn*+ cells were primarily located in the tree shrew SGS, with 90% of them co-expressing *Gad1*, confirming that they are inhibitory neurons (Figure 6I-J, S6L). In contrast, in mouse midbrain sections, *Postn* expression was detected in the Edinger-Westphal nucleus (EW) but not in the sSC, consistent with our snRNA-seq analysis (Figure 6K). Overall, our results identify an inhibitory neuron subtype in the tree shrew sSC which is absent from the mouse and provides insights into the cellular and molecular basis for its evolution.

### Conserved and divergent molecular features of the mouse, tree shrew, and human SC

Finally, we investigated the conserved and divergent molecular features of the human SC. We obtained human SC neuron data from a published snRNA-seq study (Siletti et al., 2023) (Figure S7A). Neurons (*SYN1*+/*RBFOX3*+) comprised 89% of the cells, which is consistent with the reported percentage in the original human SC dataset (Figure S7B). Integrating neurons from the human SC with our tree shrew and mouse datasets (hereafter referred to as HTM dataset) resulted in 31 clusters, annotated as HTM01 to HTM31 (Figure 7A). Three human-specific neuronal subtypes, HTM01-HTM03, were excluded from further analysis since their expression of homologous neocortical genes suggests they likely originated from the neocortex, not the SC (Figure S7C-E). The remaining 28 neuronal clusters contained cells from all three species in varying abundance, largely consistent with our previous observations in the TM dataset (Figure S7C). Importantly, neurons from the same clusters in the TM dataset consistently mapped together to corresponding clusters in the HTM dataset, indicating conserved neuronal composition across the mouse, tree shrew, and human SC (Figure 7B). As we observed when comparing tree shrew and mouse sSC neurons, cluster markers were associated with functions such as cell-substrate adhesion and neuropeptide signaling pathway and showed a higher likelihood of divergence across species (Figure 7C-D). These findings were further confirmed by examining markers of two excitatory and two inhibitory neuron subtypes (Figure S7F-G).

**Figure 7.**
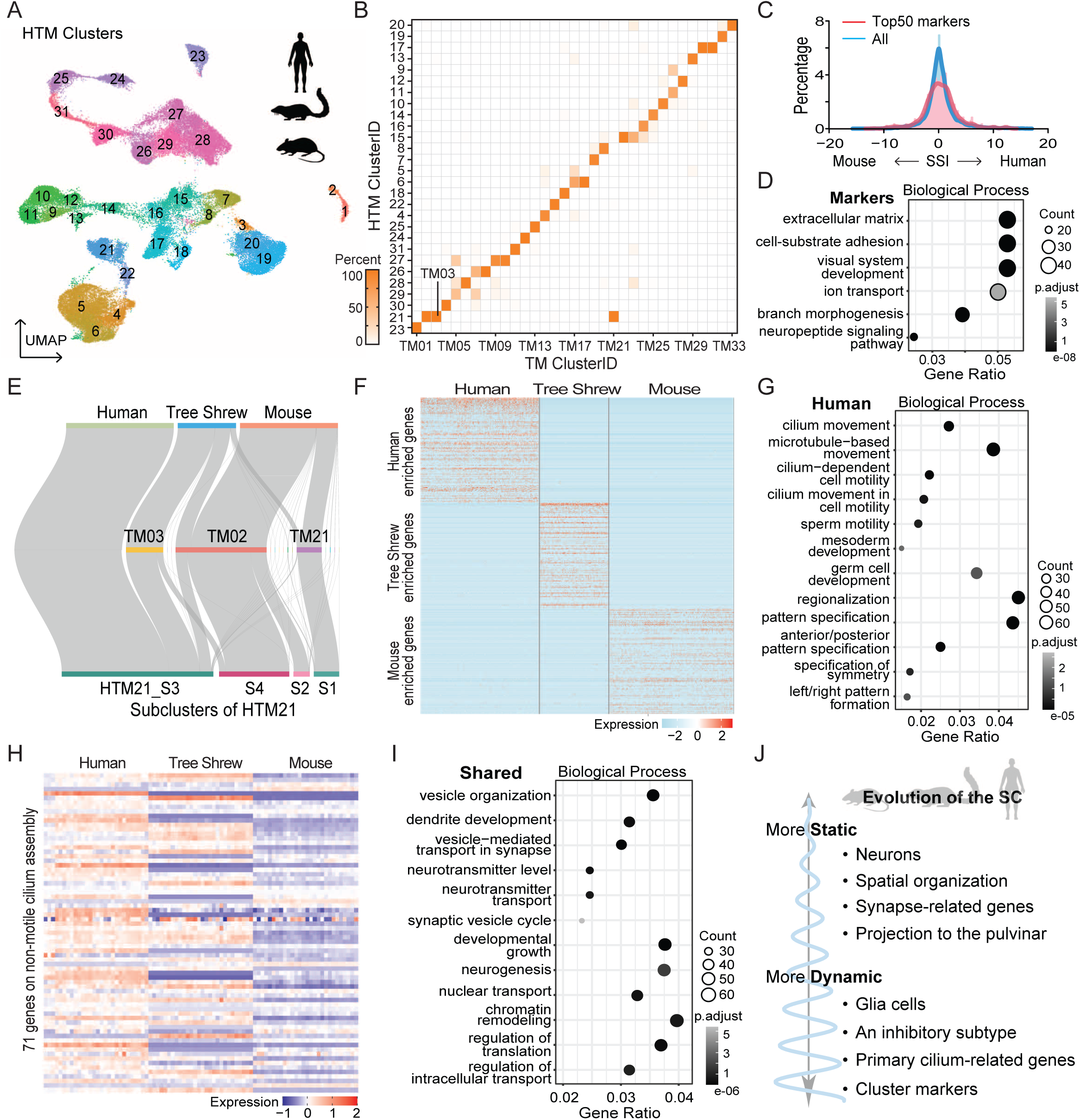
Conserved and divergent molecular features of the mouse, tree shrew, and human SC. A. UMAP plot of integrated neurons from the SC of mice, tree shrews, and humans. B. Heatmap showing the percentage of neurons from each subtype in the two-species integrated dataset (TM – tree shrew and mice) mapped to clusters in the three-species integrated dataset (HTM). C. Sankey plot illustrating the mapping of human neurons within cluster HTM21 from the HTM.integrated dataset, along with TM.integrated clusters, to the subclusters of HTM21. D. Heatmap showing the top 200 genes enriched in each species. E. GO enrichment analysis on the top 1500 human-enriched genes. F. Heatmap showing the expression of 71 primary cilium assembly-related genes across neurons in the HTM.integrated dataset. G. GO enrichment analysis on the top 1500 enriched genes shared by the three species. H. Percentage distribution of SSI between mice and humans for all genes and the cluster markers (top 50 for each cluster). I. GO enrichment analysis on the top 50 cluster markers J. Schematic summarizing the conserved and divergent molecular features of the mouse, tree shrew, and human SC.

TM03, the unique cluster in the tree shrew SC, along with TM02 and TM21, merged into HTM21, suggesting the presence of subclusters within this HTM cluster. Therefore, re-clustering only the HTM21 cells showed that almost all human HTM21 cells and most tree shrew TM03 cells clustered together, suggesting that a unique SC neuron subtype exists in both human and tree shrew SC but not in mouse SC (Figure 7E). In contrast, other subclusters of HTM21 contained almost no human SC neurons but were correspondingly mapped with neurons from TM02 and TM21, implying the absence of these two clusters in the human SC. Additionally, given the close molecular relatedness among HTM21 subclusters, the proportional differences in neurons across the three species support the possibility that these subclusters diverged from a common cellular ancestor during evolution, with human-and tree shrew-specific neuron subtype becoming predominant at the expense of other close subtypes enriched in mice.

Finally, we performed GO analysis on genes enriched in human SC relative to tree shrew and mouse SC (Figure 7F). Notably, as we previously observed of tree shrew-enriched genes, cilia-related functions were among the top GO terms for the human-enriched genes (Figure 7G). Primary cilium assembly-related genes showed higher expression in the human SC than in tree shrews and mice (Figure 7H). Other top GO terms for the human-enriched genes were related to development and organization, such as germ cell development, mesoderm development, regionalization, anterior/posterior pattern specification, and left/right pattern formation (Figure 7G). On the other hand, GO analysis of the genes shared by all three species revealed functions related to synaptic processes, such as vesicle organization, vesicle-mediated synaptic transport, and neurotransmitter regulation (Figure 7I). These results suggest that, during the evolution of sSC neurons, genes related to cilia function and developmental patterning were highly dynamic, whereas those related to synaptic function were stable. In summary, our results demonstrate that the conserved and divergent features between mice and tree shrews are consistently present in humans, offering solid insights into SC evolution at the molecular and cellular levels.

## DISCUSSION

The SC is often described as a “conserved” midbrain structure. This is based on the many anatomical features that are consistently observed in the SC (or optic tectum) across various vertebrates, including its layered organization, input/output connectivity patterns, topographic maps, and morphologically defined neuron types (Basso et al., 2021; Cang et al., 2024; Cang and Feldheim, 2013; Liu et al., 2022). Whether the molecular features and organization of the SC are conserved across mammalian species has remained unknown until now. Here, we have performed the first comparative transcriptomics study of the SC to fill this important knowledge gap and revealed molecular and cell-type divergence in a brain region which is otherwise highly conserved.

Our study yields novel insights into the molecular and cellular evolution of the nervous system. While most neuron subtypes are largely conserved across mouse, tree shrew, and human SC, some of their genes vary significantly in expression across these species. Notably, genes associated with ciliary function are among the least conserved in SC neurons across species, whereas synapse-related genes are among the most conserved. Consistent with the latter, we find that the synaptic circuitry between SC and pulvinar is similar across species – one neuronal subtype, marked by expression of the *Cbln2* gene, accounts for most SC projections to pulvinar in mouse and tree shrew. Our study also reveals how the SC has diversified with evolution. We identify a subtype of SC neuron present only in tree shrews and humans and a closely related subtype which is only in mice and tree shrews, which may represent “sister cell types” (Arendt et al., 2016). Together, our results suggest that the SC has evolved by (1) molecularly specializing cell types with otherwise conserved identities and circuitry, (2) diverging a subset of inhibitory neurons, and (3) enhancing the role of the primary cilium in neuronal signaling.

In some ways our findings align with previous comparative studies of neocortical cell types. For instance, neocortical neurons are far more conserved across species than neocortical glia (Berto et al., 2019; Jorstad et al., 2023b; Pembroke et al., 2021). Similarly, we observed that SC neurons were also more alike across species than SC glia. In addition, the human neocortex contains higher ratios of glia to neurons than the non-human primate neocortex (Sherwood et al., 2006), which we also observed in tree shrew sSC relative to mouse sSC. This suggests that, across brain regions, glia may have adapted to a greater extent than neurons to support species-specific functions.

Our study also identified other evolutionary similarities between neocortex and SC at the level of neuronal subtypes. Most neocortical neuron subtypes are homologous across mammals (Bakken et al., 2021; Hodge et al., 2019) and even vertebrates (Hain et al., 2022), with a few exceptions for species-specific cell types (Boldog et al., 2018). Likewise, we found that almost all neuron subtypes were conserved between mouse, tree shrew, and human SC. The two subtypes that were not conserved were highly similar, raising the possibility that they are ‘sister cell types’, derived from a common cell-type ancestor (Arendt et al., 2016). Comparative transcriptomic studies of the neocortex have also found species-specific neuronal subtypes (Boldog et al., 2018; Krienen et al., 2020). Interestingly, like the inhibitory neuron subtype we identified in human and tree shrew SC but not mouse SC, the species-specific neuron subtypes in human neocortex are also distinguished from other subtypes by their expression synapse-related genes (Berg et al., 2021). This raises the possibility that neuronal subtypes may diverge during evolution to form novel synaptic circuits (Kebschull et al., 2020). Another way in which the primate and mouse neocortex differs is in its layering of neuronal subtypes (Chen et al., 2023; Hodge et al., 2019). Likewise, we observed differences in the distribution of some neuronal subtypes between mouse and tree shrew SC (e.g., cluster TM13, or *Sntb1*+ neurons), though other neuronal subtypes showed a more conserved anatomical distribution (e.g., cluster TM01, or *Cbln2*+ neurons). Finally, genes with neuronal subtype-specific expression in the neocortex tended to also be specific to one species (Sousa et al., 2017), which we found was true for SC too and suggests that, across brain regions, expression of individual genes can be subject to evolution at the cell-type level.

Our study also revealed key evolutionary differences between the neocortex and SC. In the neocortex, the proportion of neurons expressing inhibitory markers increases from mouse to marmoset to human -(Bakken et al., 2021; Network, 2021) but see also (Jorstad et al., 2023a). In contrast, our study finds that the ratio of inhibitory to excitatory neurons is relatively stable across mouse, tree shrew, and human SC. This likely reflects regional differences in the importance of excitatory-inhibitory balance across mammals.

Across mammalian species, neocortical neurons differ highly in their expression of genes related to synapse function (Hodge et al., 2019; Jorstad et al., 2023b). This is true even when comparing neocortical neurons of humans and non-human primates (Ma et al., 2022; Suresh et al., 2023). The neocortex also differs across mammalian species in terms of its microcircuitry (DeFelipe et al., 2002; Peng et al., 2024). Such differences in synaptic circuitry likely require changes in neuronal expression of synaptogenic and synapse adhesion genes. On the other hand, conservation of synaptic circuitry in other regions or cell types may require similar conservation of synapse related genes. Our study found that, in contrast to the neocortex, synapse-related genes are highly conserved across mouse, tree shrew, and human SC neurons. Consistent with this, we also found that the molecular identity of pulvinar-projecting SC neurons is highly conserved between mice and tree shrews. Thus, in contrast with the neocortex, the SC exemplifies co-conservation of synaptic circuitry and gene expression.

Importantly, our study provides a foundation for future comparative studies of SC neuron function. For instance, our results demonstrate that molecular markers for cell types and neuron subtypes vary across species to greater extent than other genes. This underscores the importance of finding conserved genetic access points for investigating homologous cell types, especially given the recent development of enhancer-driven viral tools for genetically manipulating cell types in different species (Graybuck et al., 2021; Mich et al., 2021). Our study also raises questions for future investigation. For instance, since synapse-related genes are conserved in the SC but not in the neocortex, how have circuits between these two regions evolved? Future studies could assess the conservation of synapse specificity transcripts in the cortico-collicular circuit across species. Another question raised by our study is, what is driving the regional and cell type divergence in gene expression in the SC? Comparative epigenetic studies could identify changes in the presence or activity of gene regulatory elements that explain how gene expression has diverged during the evolution of this brain region.

## RESOURCE AVAILABILITY

### Lead contact

Further information and requests for experimental data should be directed to and be fulfilled by the lead contact, John N. Campbell.

### Materials availability

This study did not generate any unique reagents or materials.

### Data and code availability

The raw and quantified snRNA-seq data are available at the Gene Expression Omnibus (GEO) repository at GEO accession number ID (TBD). The code used to process and analyze the snRNA-seq data is available at Zenodo at this link: LINK (TBD). The clustered tree shrew and tree shrew/mouse integrated (TM) datasets are available for user-friendly visualization and exploration through the Broad Single Cell Portal at these links: tree shrew, LINK (TBD); tree shrew/mouse (TM dataset), LINK (TBD). The R data files (.RDS) of each of these datasets can be downloaded directly from the corresponding Broad Single Cell Portal site. All other raw and processed data are available from the corresponding authors upon request.

## ACKNOWLEDGMENTS

We thank Maosen Ye and YongGang Yao for assisting with tree shrew genome annotation and for sharing homologous gene list between mice and tree shrews. We thank Sten Linnarsson’s lab for releasing the human snRNA-seq dataset ahead of publication. Cell sorting and cytometry was performed by the University of Virginia Flow Cytometry Core Facility (RRID:SCR_017829), which is partially supported by a National Cancer Center award (P30 CA044579). Sequencing on the Illumina Next-Seq platform was performed by the Genomics Core of the Biology Department at University of Virginia and by the Genome Analysis and Technology Core of University of Virginia’s School of Medicine (RRID:SCR_018883). This work was supported by the American Diabetes Association Pathway to Stop Diabetes Initiator Award 1-18-INI-14 and US NIH grant HL153916 to J.N.C. and by US NIH grants (EY026286 and EY020950) and Jefferson Scholars Foundation to J.C..

## AUTHOR CONTRIBUTIONS

Y.L., J.N.C., and J.C. designed the experiments; Y.L. performed snRNA-seq experiments and data analysis with assistance from J.N.C; Y.L., J.A.M., and C.C. performed RNAscope experiments and data analysis; Y.L., L.Y., A.K., and A.E. performed retrograde tracer injection into the mouse and tree shrew pulvinar; E.L.S. dissected the tree shrew sSC for the sequencing experiments; Y.L., J.N.C., and J.C. wrote the manuscript with input and/or approval from all authors.

## DECLARATION OF INTERESTS

The authors declare no competing interests.

## DECLARATION OF GENERATIVE AI AND AI-ASSISTED TECHNOLOGIES

During the preparation of this work, the first author used ChatGPT to edit the language in Results, Methods, and Figure Legend in the initial draft, which was then substantially edited in multiple rounds by other authors without any AI assistance. ChatGPT was not used at all for other sections of the manuscript. The authors take full responsibility for the content of the publication.

---

**Figure S1.**
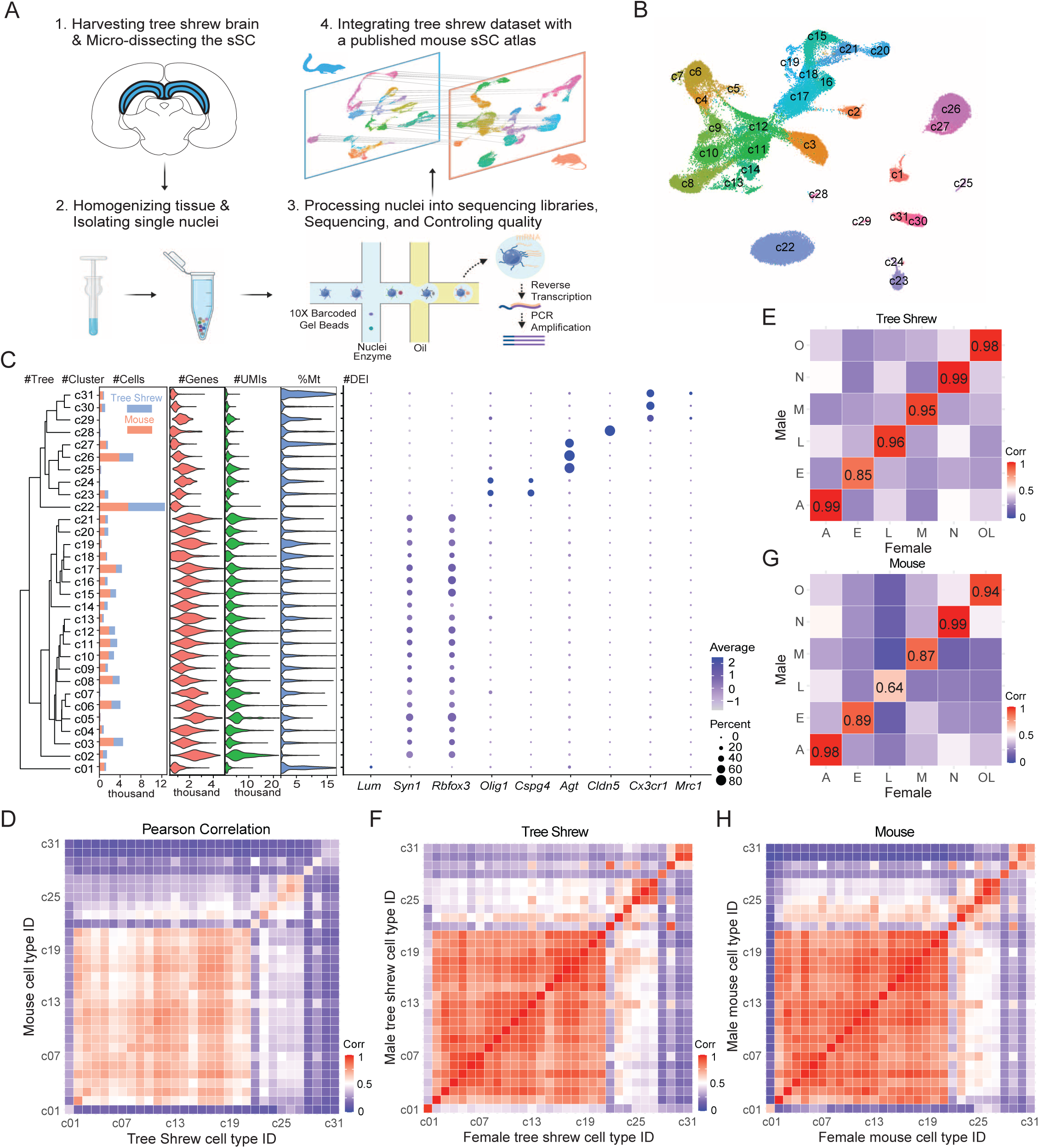
Neurons are the most conserved cell type in the mouse and tree shrew superficial superior colliculus (sSC) A. Schematic of single-nuclei RNA sequencing (snRNA-seq) workflow on the tree shrew sSC and its integration with a published mouse sSC atlas (Liu et al., 2023). B. UMAP (uniform manifold approximation and projection) plot of integrated cells from the mouse and tree shrew sSC (hereafter referred to as TM.integrated dataset), colored by cluster. Each dot represents a cell. C. Dendrogram displaying the transcriptomic relatedness of 31 cell types, followed by cluster names, cell counts per type in mice and tree shrews, the number of detected genes, transcripts, and percentage of mitochondrial genes per subtype, and a plot illustrating average expression and the percentage of cells expressing the marker genes for each cell type. Note that tree shrew cell counts overlay mouse ones. D. Pearson correlation analysis of gene expression across 31 clusters between the tree shrew and mouse. E. Pearson correlation analysis of gene expression across 6 cell types between the female and male tree shrews. F. Pearson correlation analysis of gene expression across 31 clusters between the female and male tree shrews G. Pearson correlation analysis of gene expression across 6 cell types between the female and male mice. H. Pearson correlation analysis of gene expression across 31 clusters between the female and male mice.

**Figure S2.**
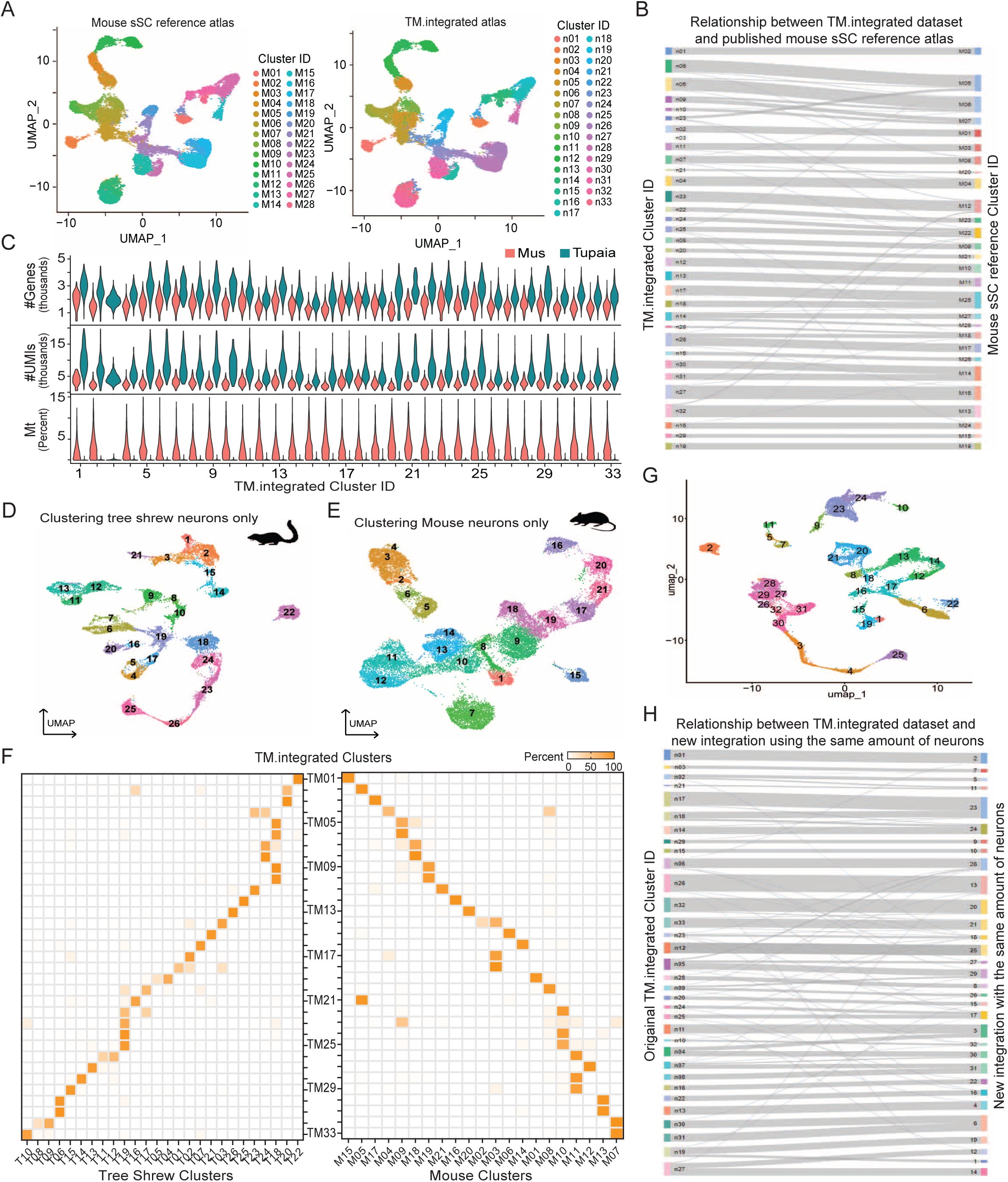
Consensus neuronal subtypes identified across clustering parameters. A. UMAP plot of the published mouse sSC reference atlas (left), alongside the visualization and annotation of the corresponding mouse neurons from the TM.integrated dataset in the same UMAP space (right). B. Sankey plot illustrating the cluster relationships between the TM.integrated dataset and the published mouse sSC reference atlas. Mouse neurons from the same clusters in the TM dataset are generally mapped together onto corresponding clusters, with some subtypes merging into a single subtype in the mouse sSC dataset. The thickness of each band represents the relative number of neurons within that cluster. C. The number of detected genes, transcripts, and percentage of mitochondrial genes in each neuronal subtype of tree shrews and mice. D. Clustering only tree shrew sSC neurons with the same parameters used in the TM.integrated dataset. E. Clustering only mouse sSC neurons with the same parameters used in the TM.integrated dataset. F. Percentage of neurons in separately identified clusters for tree shrews (left) and mice (right), mapped to the neuronal subtypes of the TM.integrated dataset. The one-to-one cluster correspondence suggests that the analysis was primarily driven by molecular profiles rather than technical artifacts. G. Clustering equal number of mouse and tree shrew neurons with the same parameters used in the TM.integrated dataset. H. Clustering relationship between TM.integrated dataset and the integration of equal sample sizes. Robust correspondence of clusters was observed between before and after downsampling, suggesting that higher number in mice do not dominate the clustering results of integration

**Figure S3.**
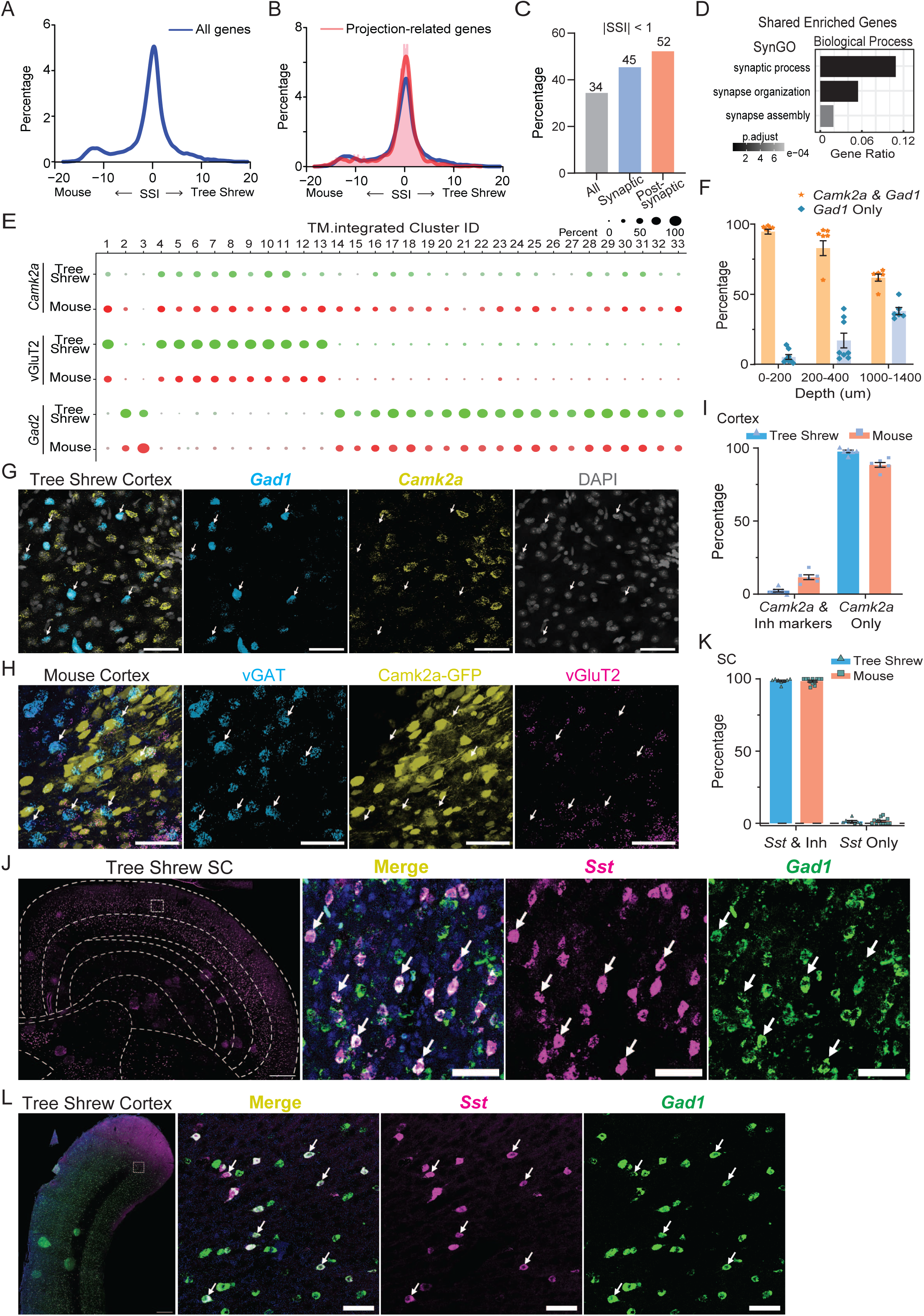
Synapse-related genes are conserved between the mouse and tree shrew sSC. A. Percentage distribution of species specificity index (SSI) for all genes. Positive x-axis values: higher gene expression in tree shrews; negative: higher in mice. B. Percentage distribution of SSI for projection-related genes. C. Percentage of genes with an absolute SSI value lower than 1 among all genes, synapse-related genes, and postsynapse-related genes. D. SynGO enrichment analysis related to biological process for the top 1600 shared genes E. Average expression and percentage of neurons expressing Camk2a, vGluT2, and Gad2 in each neuronal subtype by species. F. Percentage of Gad1+ cells co-expressing Camk2a across the depth of the tree shrew SC. Mean ± SEM. n = 8 ROIs; n = 2 animals. G. FISH of Gad1 and Camk2a expression in the tree shrew cortex. Scale bar, 50 μm. H. FISH of vGAT and vGluT2 expression in the mouse cortex, with neurons labeled by AAV-Camk2a-GFP. Scale bar, 50 μm. I. Percentage of Camk2a+ cells co-localized with inhibitory markers in mouse and tree shrew cortex. Mean ± SEM. n = 6 ROIs; n = 3 animals each. J. FISH of Sst and Gad1 expression in the tree shrew SC. Scale bar, 500 μm (left), 50 μm (right). K. Percentage of Sst+ cells co-expressing inhibitory markers in the tree shrew and mouse SC. Mean ± SEM. n = 8 ROIs in tree shrew and 12 ROIs in mice; n = 2 tree shrews and 3 mice. L. FISH of Sst and Gad1 expression in the tree shrew cortex. Scale bar, 500 μm (left), 50 μm (right). B

**Figure S4.**
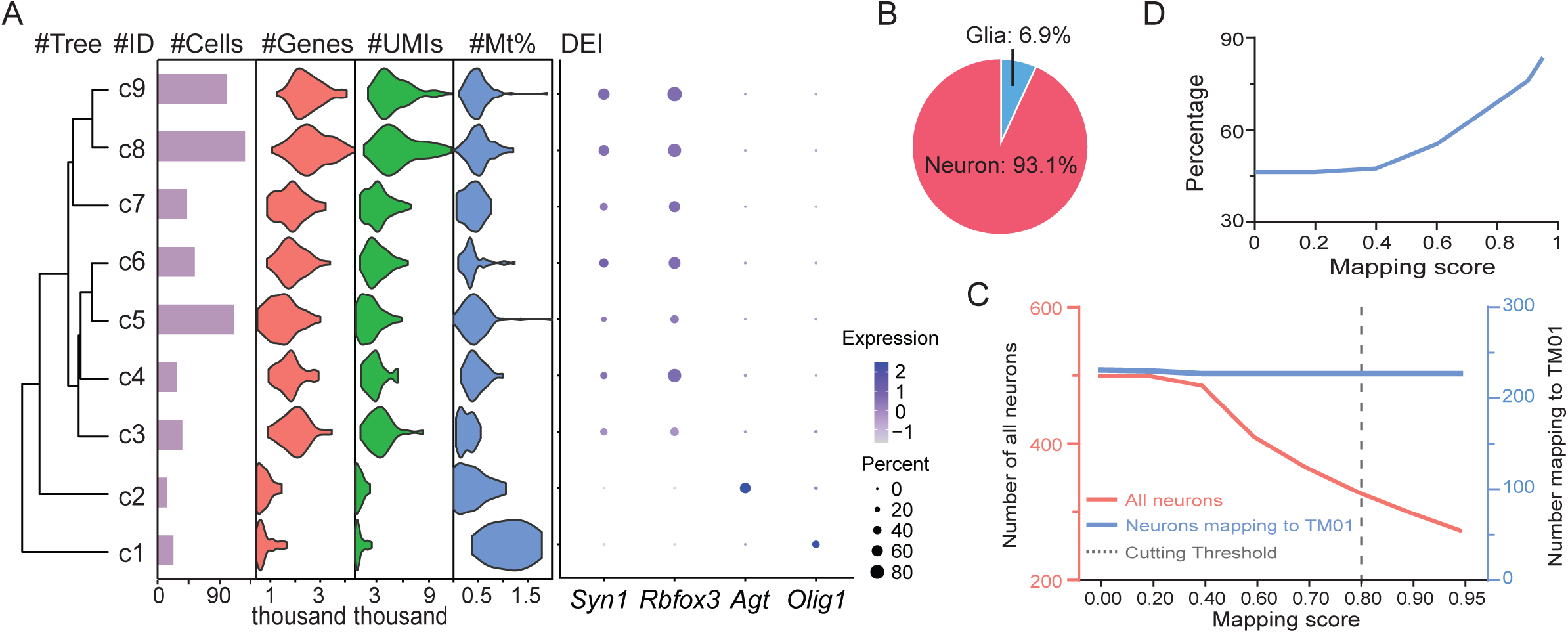
Pulvinar-projecting neurons in mice map to cluster TM01. A. Dendrogram illustrating the transcriptomic relatedness of 9 cell types, followed by cluster names, cell counts per cluster, the number of detected genes and transcripts, the percentage of mitochondrial genes per cluster, and a plot illustrating average expression and percentage of cells expressing the cell type marker genes. B. Percentage of neurons and glial cells in the profiled population. C. Total number of remaining neurons and those mapped to Clu.TM01 after filtering by the confidence score. D. Percentage of neurons mapped to Clu.TM01 out of the total number of neurons after filtering by the confidence score

**Figure S5.**
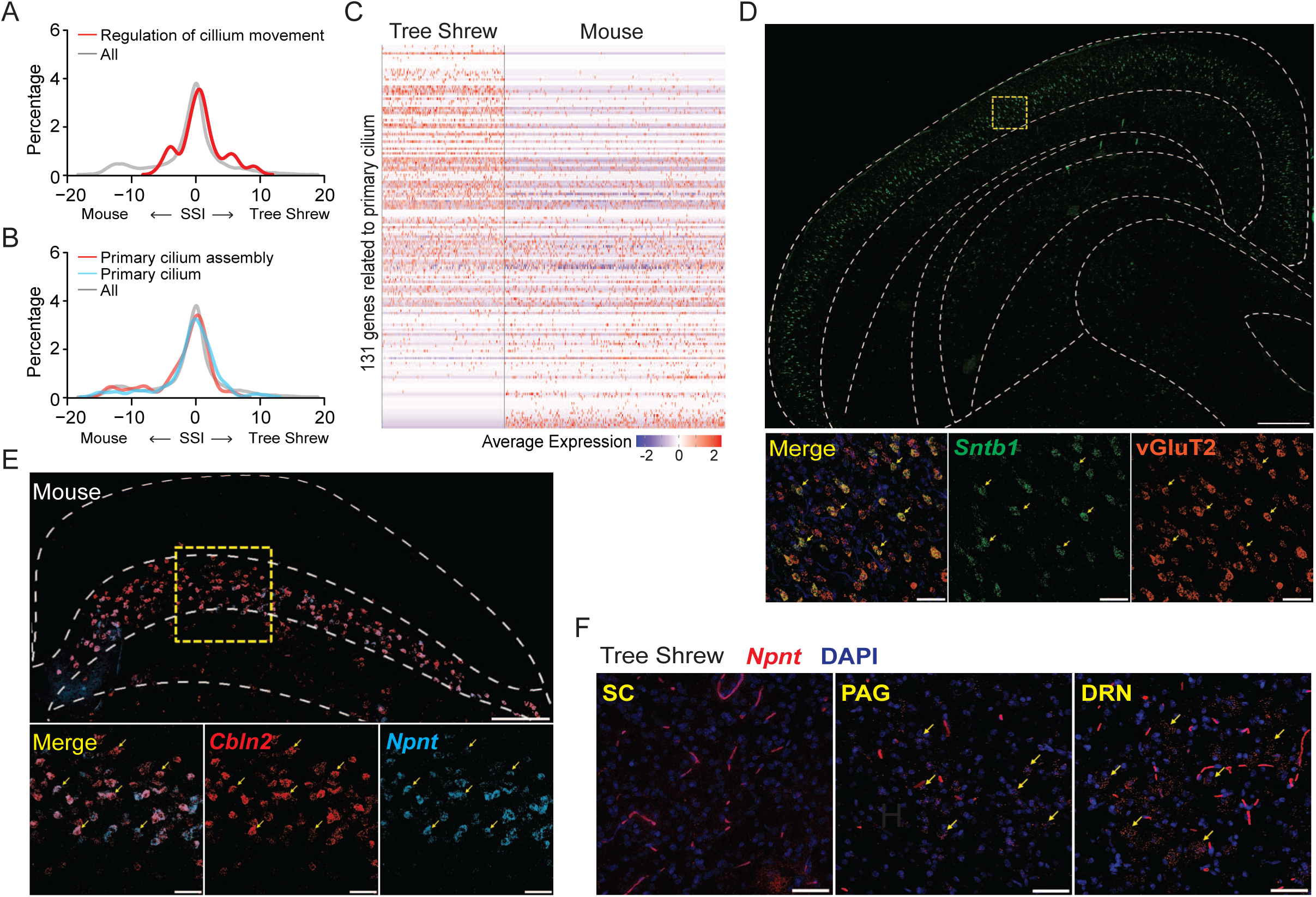
Divergent gene expression between tree shrew and mouse sSC neurons. A. Percentage distribution of SSI for all genes and genes associated with regulation of cilium movement. B. Percentage distribution of SSI for genes associated with primary cilium and primary cilium assembly. C. Heatmap showing expression level of 131 primary cilium-related genes across neurons in the dataset. D. FISH of Sntb1 and vGluT2 in the tree shrew SC. Scale bar, 500 μm (top), 50 μm (bottom). E. FISH of Cbln2 and Npnt in the mouse SC. Scale bar, 200 μm (top), 50 μm (bottom). F. FISH of Npnt in the tree shrew midbrain. Npnt expression was detected in the periaqueductal gray (PAG) and the dorsal raphe nucleus (DRN), but not in the SC. Scale bar, 50 μm.

**Figure S6.**
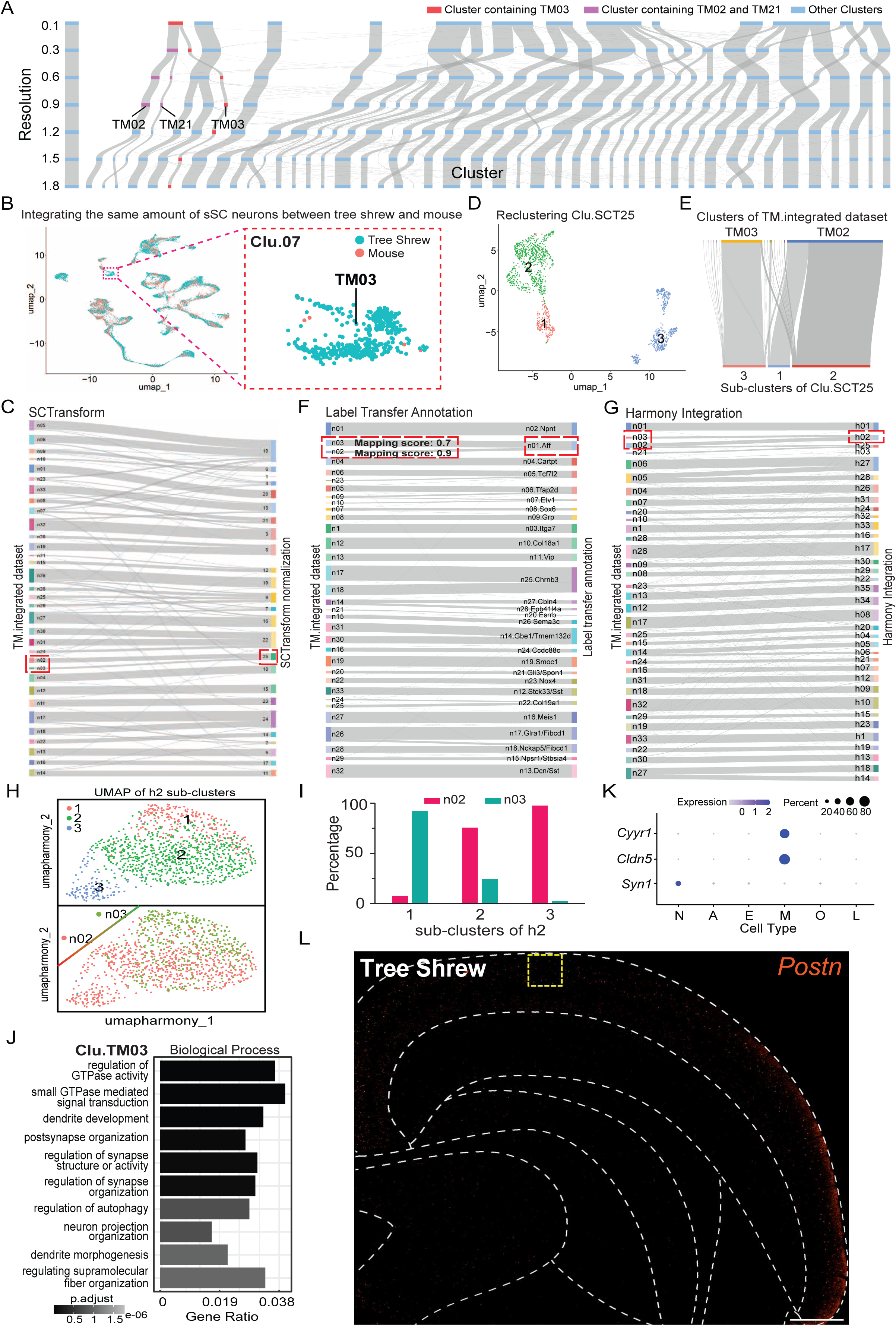
An inhibitory neuron subtype is detected in the tree shrew but not mouse sSC. A. Clustering relationship across various clustering resolutions. TM03 appeared at a resolution of 0.3, which is lower than the recommended value for optimal clustering, and remained unique at higher resolutions, suggesting its stable identification across resolution. The cluster containing TM03 cells is labeled in red, whereas clusters with close transcriptomic relatedness, TM02 and TM21, are labeled in magenta. B. UMAP plot showing clustering of equal numbers of mouse (red) and tree shrew (cyan) neurons. Neurons from TM03 are grouped together as Clu.07, shown at a higher magnification on the right. C. Clustering relationship between the TM.integrated dataset and the integration using SCTransform normalization. Neurons from TM03, alongside those from TM02, merged into SCT25, which was highlighted in a red frame. D. UMAP plot showing subtypes within Clu.SCT25. E. Clustering relationship between TM.integrated dataset and the subtypes of Clu.SCT25, with almost all neurons from TM03 mapped to a distinct subtype unique to tree shrews. F. Clustering relationship between the TM.integrated dataset and the integration using the label transfer annotation method. Neurons from TM03 and TM02 were mapped to n01.Aff, highlighted in a red frame. The average mapping accuracy for TM03 (0.7 out of 1) was lower compared to glial cells (0.8), whereas TM02 was higher (0.9), suggesting that TM03 might represent a distinct cluster but is being forced to fit into the reference atlas. G. Clustering relationship between the TM.integrated dataset and the integration using the Harmony integrative method. Neurons from TM03 and TM02 were mapped to cluster h02. H. UMAP plot showing subclusters within h02 and their relationship with TM02 and TM03. I. Percentage of neurons from TM02 and TM03 mapped to each subcluster of h02. One subcluster primarily contains neurons from TM03 and another predominantly consists of neurons from TM02. J. GO enrichment analysis of marker genes from TM03. K. Average expression and percentage of cells expressing *Syn1* (marker for neurons), *Cldn5* (marker for endothelial cells), and *Cyyr1* across cell types. Though *Cyyr1* expression was enriched in TM03, its relative expression in neurons was much lower than endothelial cells, suggesting that it is not a good marker for FISH. L. FISH of Postn expression in the tree shrew SC. Scale bar, 500 μm.

**Figure S7.**
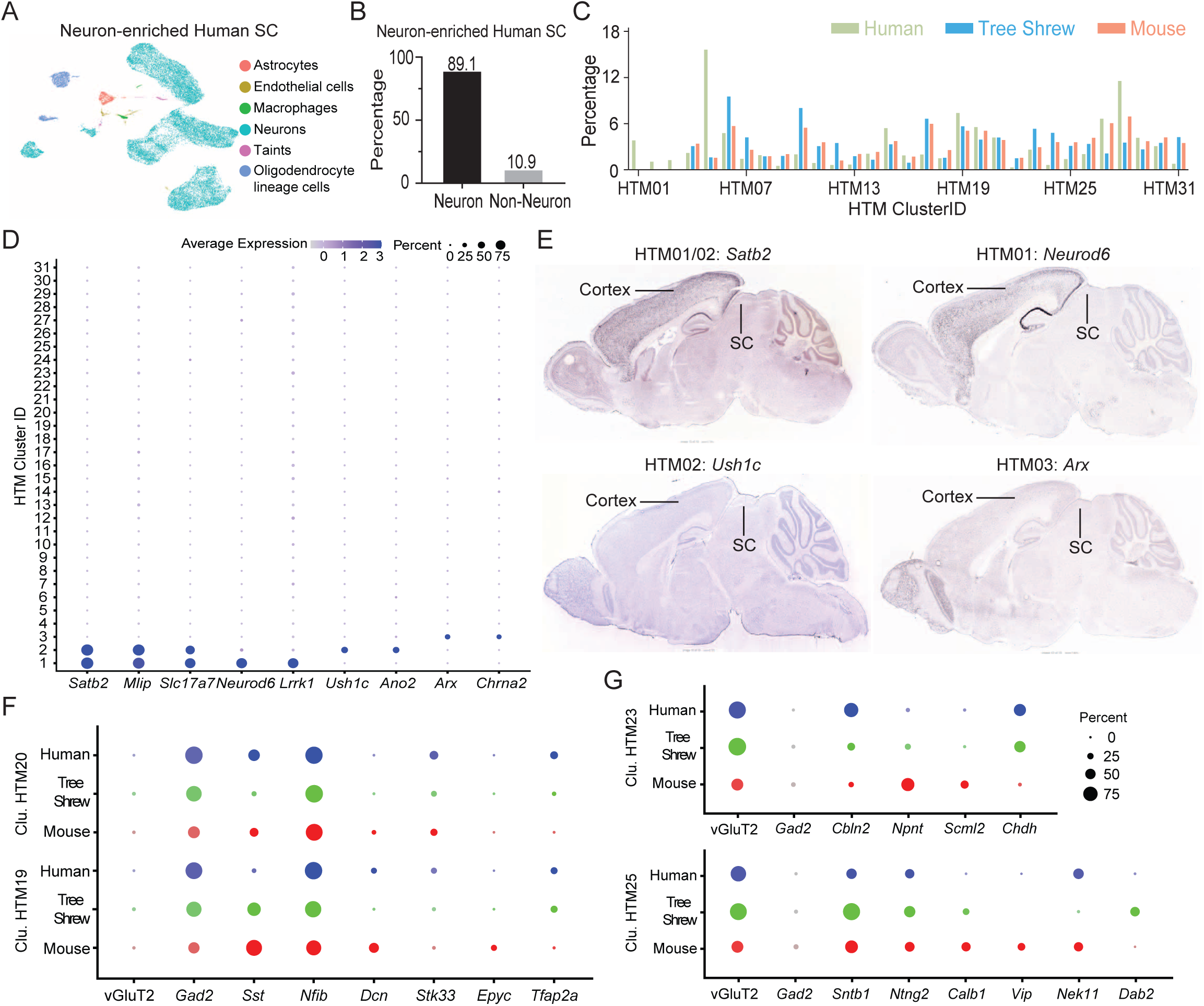
Shared and divergent features between mouse, tree shrew, and human SC. A. UMAP plot showing clustering of neuron-enriched human SC snRNA-seq dataset, colored by cell type. B. Percentage of neurons and non-neurons identified in our analysis of the neuron-enriched human SC dataset. C. Percentage of neurons in each neuronal subtype by each species in the three-species integrated dataset (HTM). D. Average expression and percentage of neurons expressing marker genes selectively enriched for cluster HTM01, HTM02, and HTM03. E. In situ hybridization images from the Allen Mouse Brain Atlas showing expression of cluster markers for HTM01, HTM02, and HTM03 in the mouse cortex, but not in the SC. F – G. Average expression and percentage of neurons expressing marker genes for two example inhibitory clusters (HTM19 and HTM20) and two example excitatory clusters (HTM23 and HTM25).

## METHODS

### Animals

A total of 9 adult male and female wild-type tree shrews, aged 6 to 16 months, were used in this study. Specifically, one male and one female tree shrew were used for snRNA-seq of the superficial superior colliculus (sSC); three for retrograde tracer injections into the pulvinar; and another four for RNA fluorescence in situ hybridization (RNA FISH). Additionally, 21 mice aged 2 to 8 months were used, including H2b-TRAP (Jackson Laboratory, Strain #029789; RRID: IMSR_JAX: 029789) and C57Bl/6J (Strain #000664; RRID: IMSR_JAX: 000664). Of these, nine H2b-TRAP mice were used for snRNA-seq of pulvinar-projecting SC neurons; six were used for RNA FISH; three for CTB injections to confirm molecular identity of the pulvinar-projection neurons; and three for injections of AAV-Camk2a-GFP.

All tree shrews and mice were kept on a 12hr light/dark cycle under standard feeding and housing conditions, with 1 to 2 tree shrews housed per cage and 2 to 5 mice housed per cage. All experimental procedures were approved by the University of Virginia Institutional Animal Care and Use Committee.

### Single-nuclei RNA-sequencing of tree shrew sSC

Tree shrews were anesthetized first with isoflurane and then with a lethal dose of euthasol. Brains were rapidly dissected, cooled in ice-cold Hibernate-A for 1.5 minutes, transferred to PBS, and prepared under a dissecting microscope. The cortex and cerebellum were removed to expose the superior colliculus (SC). The sSC was visualized coronally, micro-dissected by knife cuts along its visually approximated anatomical borders, and processed into a single-nucleus suspension using a published detergent-mechanical lysis protocol (Matson et al., 2018). Briefly, the sSC was homogenized in a lysis buffer, and cell nuclei were purified by density gradient centrifugation. After resuspension, nuclei were labeled with propidium iodide (Propidium Iodide Ready Flow™ Reagent, Invitrogen, catalog # R37169) and then counted with a CellDrop automated cell counter (DeNovix, Delaware, USA).

We used the 10X Genomics Chromium Next GEM 3’ kit v3.1 according to the manufacturer’s protocol (user guide revision D) to encapsulate each nucleus’s poly-adenylated RNA and then process the poly-adenylated RNA into cDNA sequencing libraries. We then sequenced the cDNA libraries by Illumina Next-Seq 2000 to an average of 29,857 mean reads per cell. We loaded approximately 16,500 nuclei per lane of the 10X chip. A total of 4 single-nuclei samples were processed across 2 10X chips (batches): samples 1 and 2 were aliquots of the same nuclei suspension from a female tree shrew and processed in parallel as batch 1; samples 3 and 4 were aliquots of the same nuclei suspension from a male tree shrew processed in parallel as batch 2. Finally, the 10X Cell Ranger pipeline (version 7.2.0) was used to demultiplex and align sequencing reads to a custom “pre-mRNA” reference genome created from the published tree shrew database (http://www.treeshrewdb.org/) to generate feature-barcode matrices for each batch of tree shrew sSC.

### Integrating snRNA-seq datasets from the mouse and tree shrew sSC

The mouse sSC feature-barcode matrices were obtained from our previous publication (Liu et al., 2023). Both the tree shrew and mouse feature-barcode matrices were analyzed using R (version 4.3.1) with the Seurat 5.0.3 software package for R (Hao et al., 2024). For the tree shrew samples, quality control was performed on both batches, filtering out genes detected in fewer than three cells and cells with fewer than 200 genes, more than 6,500 genes, or over 21,000 transcripts. We filtered the mouse dataset as previously described (Liu et al., 2023) - in general, filtering out genes detected in fewer than three cells and cells with fewer than 200 genes, more than 3,500 genes, or over 7,200 transcripts. To place cells from the two species into the same gene expression space, the tree shrew and mouse matrices were filtered so that only orthologous genes were used for dataset integration, based on the orthologous gene list provided by Dr. Mao-sen Ye from the Kunming Institute of Zoology.

Next, we integrated the tree shrew and mouse matrices to correct for technical variance including batch effects. In brief, we log-normalized the data; selected 2,000 most variable genes for each matrix; found anchor correspondences across matrices with canonical correlation analysis (CCA); integrated the matrices using the IntegrateData() function in Seurat; scaled each gene; performed Principal Component Analysis (PCA) to linearly reduce the dimensionality of the highly variable gene set; clustered the cells using the Louvain algorithm, based on Euclidean distance in the PCA space comprising the first 10 PCs and with a resolution value of 1; reordered the cluster ID using the BuildClusterTree() function; and performed non-linear dimensionality reduction by Uniform Manifold Approximation and Projection (UMAP). Cell types were annotated according to their expression of classical cell type marker genes: neurons (*Syn1*+, *Rbfox3*+); oligodendrocyte lineage cells (*Olig1*+, *Cspg4*+); astrocytes (*Agt*+); endothelial cells (*Cldn5*+); macrophages (*Cx3cr1*+, *Mrc1*+); and leptomeningeal cells (*Lum*+; Figure 1A, S1C). Clusters potentially representing cell doublets were identified based on cluster-and cell-level co-expression of these marker genes. After removing the suspect clusters, the remaining cells were re-clustered in two additional rounds to remove remaining potential doublets, following the steps of log normalization, feature selection, anchor-based integration, PCA, and clustering (top 10 PCs, resolution set to 1).

Clusters with relatively high expression of the neuronal marker genes *Syn1* and *Rbfox3*, but little to no expression of other cell type marker genes were annotated as neurons, then subset and re-clustered for another two additional rounds to remove suspected cell doublets. In the first round of neuronal re-clustering, the top 15 PCs and a resolution value of 1 were used, while the second round used the top 14 PCs with a resolution value of 0.9. A total of 33 neuronal clusters were identified from the tree shrew and mouse integrated dataset (hereafter referred to as TM dataset). Genes differentially expressed across clusters were identified using the Wilcoxon Rank Sum test and filtered to include only those detected in at least 10% of cells in the cluster. Candidate marker genes were selected based on their expression least detected by other clusters and highly expressed in the given clusters.

### Clustering relationships across various parameters and integrative methods

To explore the relationship between neuronal subtypes in the TM dataset and mouse sSC datasets, individual mouse neurons were matched between these datasets according to 10X cell-specific barcodes. The corresponding neurons were visualized using givsSankey() function based on ‘googleVis’ package (Figure S2A-B). To reduce the technical bias, the clustering analysis was also performed independently for the mouse or the tree shrew neurons while using the orthologous genes and keeping the same PCs and resolution as in the TM dataset. Corresponding cells of the integrated and species-specific datasets were also visualized with the Sankey plot (Figure S2C-F).

Given that the TM dataset includes more neurons from mice (27,497) than from tree shrews (15,443), we assessed whether the larger sample size might influence the clustering results of the integrated data. An equivalent number of mouse neurons were randomly selected and integrated with tree shrew neurons using the same parameters as in the TM dataset (Figure S2G-H). Additionally, resolution parameters were tested varying from 0.1 to 1.8 for clustering with the same 14 PCs, as a high resolution might lead to overfitting (Figure S6A). Moreover, while maintaining the same PCs and resolution, different normalization and integration methods were tested, such as SCTansform normalization with the SCTransform() function, Harmony with the IntegrateLayers() function, label transfer annotation with the TransferData() function, and reciprocal principal component reduction in the FindIntegrationAnchors() function (Figure S6C-G).

### Integrating human, tree shrew, and mouse SC datasets

The human SC dataset was obtained from the Human Brain Cell Atlas v1.0 via the CELLxGENE platform (https://github.com/linnarsson-lab/adult-human-brain). In brief, this dataset was generated in a previous study from three adult male donors using the 10X Chromium Next GEM 3’ reagent kit v3 for sequencing library preparation and sequenced on an Illumina NovaSeq platform. Sequencing reads were mapped to the human reference genome GRCh38.p13 (Gencode V35) to generate a feature-barcode matrix using Cell Ranger version 4.0.0. After obtaining this human SC snRNA-seq dataset, we first converted its features from Ensembl IDs to gene symbols using the "org.Hs.eg.db" package in R, and then further converted it into mouse orthologous genes using the convert_human_to_mouse_symbols() function in the "nichenetr" package (Browaeys et al., 2020).

Next, we conducted quality control on the human SC dataset based on its quality metric distribution. Cells with more than 8,000 detected genes, over 60,000 transcripts, or a mitochondrial gene fraction greater than 0.3 were filtered out. Subsequently, a similar clustering approach was used to identify cell types within the entire human SC dataset, including log normalization, feature selection, data scaling, PCA, clustering (using the top 9 PCs and a resolution of 1), and identification of marker gene expression. Clusters with relatively high expression of the neuronal marker genes *Syn1* and *Rbfox3* were subset and re-clustered to further exclude potential cell doublets, using the top 6 PCs and a resolution of 1.

The human neuronal feature-barcode matrix was then integrated into the TM dataset. This process involved log normalization, feature selection, anchor identification using CCA, matrix integration, scaling, PCA, and clustering (using the top 11 PCs with a resolution of 1). As a result, we identified 31 neuronal subtypes (hereafter referred to as HTM dataset). Candidate marker genes were selected based on their expression in more than 10% of cells of the given cluster and little to no detection in other clusters.

### Analysis of conserved and divergent genes across clusters and species

To compare gene expression across species, we conducted a differentially enriched gene (DEG) analysis using Seurat’s FindMarker() function with the Wilcoxon Rank Sum test. To ensure all genes were included, we adjusted the default settings by setting the minimum fraction of detected genes in either species to 0 and the log2 fold change threshold to 0. Due to minor differences between the list of orthologous genes in mice and tree shrews compared to mice and humans, only orthologous genes were included in this analysis. Specifically, orthologous genes between tree shrews and mice were used for the TM dataset, whereas orthologous genes across the three species were used for the HTM dataset. Additionally, we tested that this minor difference did not impact the integration results.

The cross-species DEG analysis yielded the log2 fold change of expression means between species (hereafter referred to as “species specificity index (SSI)”), the percentage of cells in which the gene was detected in each species, and a p-value adjusted according to Bonferroni correction. Positive SSI values indicate enrichment in tree shrews, whereas negative SSI values indicate enrichment in mice. To identify species-specific genes in the TM dataset, we selected 1,600 genes with SSI values higher than 4 (tree shrew-enriched) and 1,600 genes with SSI values lower than -9 (mouse-enriched). Additionally, we selected 1,600 shared genes with absolute SSI values lower than 0.27 (most conserved genes between mice and tree shrews). On the other hand, to identify human-specific genes or shared genes shared by the three species in the HTM dataset, we performed DEG analysis comparing gene expression in human neurons with those in tree shrew and mouse neurons. The top 1,500 genes with the highest SSI values were selected as human-specific genes, whereas those with the lowest absolute SSI values were identified as shared genes.

To predict the functions of conserved and species-enriched genes, we performed Gene Ontology (GO) enrichment analysis using the enrichGO() function from the “clusterProfiler” and “org.Mm.eg.db” packages in R (Yu et al., 2012). Moreover, to confirm that the shared genes were related to synapse function, we conducted a synGO analysis which is designed specifically for synapse function (https://www.syngoportal.org/) (Koopmans et al., 2019). On the other hand, we examined distribution of SSI values on selected GO terms between tree shrews and mice or across three species, including: synapse (GO:0045202); postsynapse organization (GO:0099173); cell projection organization (GO:0030030); cilium movement (GO:0003341); secretory granule (GO:0030141); non-motile cilium assembly (GO:1905515); non-motile cilium (GO:0097730); and positive regulation of cilium movement (GO:0003353).

To calculate the log2 fold change for all genes across clusters (cluster Log2FC) in the TM dataset and HTM dataset, we used the FindAllMarkers() function in Seurat. We then reordered the genes in the resulting sheet based on their Log2FC values, retained the first occurrence of duplicated genes with the highest values, and removed the other duplicates. In addition, we identified the top 50 cluster markers for each subtype based on the difference in the percentage of neurons in which expression of the gene was detected, inside vs. outside the selected cluster. This identified a total of 982 cluster markers across all clusters.

### snRNA-seq of pulvinar-projecting neurons in mouse SC

Four weeks after pENN.AAV.hSyn.Cre.WPRE.hGH (Addgene, Catalog # 105553-AAVrg) was injected into the pulvinar of H2B-TRAP mice, their brains were harvested following rapid decapitation to minimize the effects of stress on gene expression. The brains were cooled in ice-cold PBS for 1.5 minutes, placed in a chilled stainless steel brain matrix, and coronally sectioned into 1mm thick slices containing the sSC (Bregma -3.0 mm to -5.0 mm). The slices were immediately immersed in ice-cold RNAprotect reagent (Qiagen) and incubated at 4°C overnight to stabilize RNA. The following day, the sSC was visualized under a fluorescent dissecting microscope, micro-dissected using knife cuts, and processed into single-nuclei suspensions using a detergent-mechanical lysis protocol. After resuspending the nuclei pellet, DRAQ5 was added, and mCherry-positive nuclei were sorted using a Becton Dickinson Influx Cell Sorter. Poly-adenylated RNA from each sorted nucleus was encapsulated and used to generate cDNA sequencing libraries with the 10X Chromium Next GEM 3’ reagent kit v3.1, following the user guide revision D protocol. cDNA libraries were then sequenced on an Illumina NextSeq 2000 platform to an average depth of 28,135 reads per cell. After that, Cell Ranger (version 5.0.0) was used to map sequence reads to a custom "pre-mRNA" reference genome, built from the mouse reference genome GRCm38.98, to generate a feature-barcode matrix.

The feature-barcode matrix was analyzed using the Seurat 5.0.3 package (Hao et al., 2024) in R (version 4.3.1). Genes detected in fewer than three cells, and cells with fewer than 200 or more than 5,000 detected genes, over 12,500 UMIs, or more than 2% mitochondrial content were filtered out. Then a similar procedure for clustering was conducted, including log normalization, feature selection, scaling, PCA, clustering with the top 9PCs and a resolution of 0.5, and detected expression of cell type markers. Clusters with enriched expression of the neuronal markers *Syn1* and *Rbfox3* were subset and re-clustered to identify neuronal subtypes (top 15 PCs and resolution at 0.3). To determine the identities of these neuronal subtypes, we used the label transfer annotation method to map them to the TM dataset, with the FindTransferAnchors() and TransferData() functions in Seurat. A confidence score at 0.8 was applied to minimize technical bias when assigning cells to the reference atlas.

### Stereotaxic injection of tracers and viral vectors

Mice were anesthetized with 2% isoflurane both before and during surgery and positioned in a stereotaxic apparatus. A small midline incision was made to expose the skull, and the skull was adjusted to ensure it was flat by aligning Bregma and Lambda (Anterior-Posterior) at the same level, as well as the left and right sides (Medial-Lateral). The injection site was marked based on the targeted coordinates relative to Bregma, and the skull at the marked location was carefully drilled. Three sets of injections were then performed with pressure injector (Drummond Nanoject II, Cat# 3000204): (1) 13.8 nl of pENN.AAV.hSyn.Cre.WPRE.hGH (Addgene, Catalog # 105553-AAVrg) was injected into the pulvinar (ML: 1.4 mm; AP: -2.3 mm; DV: -2.55 mm) of H2B-TRAP mice to label pulvinar-projecting neurons; (2) three injections of 46 nL of pAAV-CaMKIIa-EGFP (Addgene, Catalog #50469-AAV9) were delivered into the cortex (AP: 0.8 mm; ML: -3.8 mm; DV: 0.7 mm) and two sites in the SC (ML: 0.8 mm; AP: -3.8 mm; DV: 1.2 & 1.7 mm) of WT mice; and (3) 45nl of Cholera Toxin Subunit B, conjugated with Alexa Fluor 555 (5 mg/mL; Invitrogen by Thermo Fisher Scientific, Catalog # C34776) was injected into the pulvinar (ML: 1.4 mm; AP: -2.3 mm; DV: -2.55 mm) of WT mice. After that, the holes in the skull were filled with wax, the scalp was closed with sutures, and mice were injected with 2% carprofen.

Tree shrews were initially anesthetized with 5% isoflurane, followed by intraperitoneal injections of pre-surgical medications including midazolam (5 mg/kg), atropine (0.3 mg/kg), and dexamethasone (1 mg/kg). Afterward, the animals were positioned in a stereotaxic apparatus with maintenance anesthesia at approximately 3% isoflurane. EKG pads were attached to the limbs to monitor heart activity, and once stable EKG signals were recorded, the animals were intubated. A similar stereotaxic procedure used for mice was then applied to the tree shrews, including scalp incision, Bregma-Lambda alignment, marking target coordinates, and drilling at the marked location. pAAV-CAG-tdTomato viral vector (Addgene, Catalog # 59462-AAVrg) was injected into three sites of the pulvinar (ML: 3 mm; AP: -4.28 mm; DV: 5.00/5.25/5.55 mm) of the tree shrew, with 200 nL delivered at each depth. The injection sites were sealed with wax, and the scalp was closed with sutures. Following the surgery, the animals were administered post-surgical medications (Buprenorphine: 0.05 mg/kg; Enrofloxacin: 5 mg/kg) and placed on a heating pad until fully recovered from anesthesia.

### Perfusion and histology

Tree shrews were initially anesthetized with isoflurane in a chamber, followed by deep anesthesia with an intraperitoneal injection of a lethal dose of Euthasol. Mice were anesthetized directly with intraperitoneal Euthasol. All animals were then perfused with PBS, followed by 4% paraformaldehyde (PFA). Brains were dissected immediately, fixed in 4% PFA overnight, dehydrated in 30% sucrose for three days, embedded in O.C.T. compound, frozen at -20 °C, and sectioned coronally at 40 μm thickness using a cryostat (Leica CM1950). The slices were finally collected and stored in cryoprotectant at -20 °C.

### RNA fluorescence in situ hybridization (FISH)

Slices removed from cryoprotectant were washed twice in phosphate-buffered saline (PBS; pH 7.2) for 5 minutes each, then incubated with hydrogen peroxide for 10 minutes within a well plate. After incubation, the slices were washed three times in PBS to remove bubbles and mounted onto slides (Fisher Cat. 12-550-15). The mouse slices were air-dried overnight in preparation for RNA FISH the next day. In contrast, slices from tree shrews underwent target retrieval before FISH. Specifically, tree shrew slices were baked in an oven at 40 °C for 30 minutes, incubated in 4% PFA at 4 °C for 15 minutes, and dehydrated sequentially in 50%, 70%, and 100% ethanol at room temperature for 5 minutes each. The slices were then air-dried for 5 minutes, incubated in distilled water at 99 °C for 10 seconds, followed by antigen retrieval solution at 99 °C for 5 minutes, rinsed in distilled water for 15 seconds at room temperature, and finally dehydrated in 100% ethanol for 3 minutes.

On the following day, RNAscope Multiplex v2 (Cat. 323100) was used to stain specific messenger RNA transcripts in the coronal sections, following the manufacturer’s protocol. Briefly, slide-mounted slices were circled with a hydrophobic barrier, rinsed with distilled water, digested with Protease IV at 40 °C for 30 minutes, and incubated with targeted probes at 40 °C for 2 hours. The slices were then treated with amplification reagents (Amp 1-3) at 40 °C. The probes used were: *Slc17a6* (vGluT2); *Slc32a1* (vGAT); *Gad1*; *Camk2a*; *Cbln2*; *Cbln4*; *Sst*; *Npnt*; *Vip*; *Sntb1*; *Postn*; and *tdTomato*. Following this, HRP-C1/C2/C3, FITC/Cy3/Cy5 dyes, and HRP blocker were applied to bind to Amp 3 and activate fluorescence. Finally, DAPI was used as a counterstain, and fluorescence images were captured using a confocal microscope (20X/0.80 Plan Apo, Zeiss LSM 800).

### RNA FISH analysis

To reveal the spatial organization of gene expression in the tree shrew and mouse superior colliculus (SC), the layering structures were outlined using the Allen Mouse Brain Atlas (Lein et al., 2007) (https://mouse.brain-map.org/) as a reference and the stereotaxic coordinates of the tree shrew brain. Based on these structural outlines and the density of detected expression of inhibitory and excitatory markers, we identified a correspondence in depth between the two species (Figure 2D). Specifically, the SO sublayer in the tree shrew SC begins at approximately 500 µm and ends around 800 µm from the surface, whereas in mice, it starts at about 300 µm and ends around 500 µm (Figure 2F, G).

To investigate the spatial distribution of inhibitory and excitatory neurons along the depth of the SC in tree shrews, we performed separate RNA FISH experiments for *Gad1* and vGluT2 on different tree shrew slices. This was necessary to prevent signal leaking between channels during confocal imaging due to their robust expression. Then a 1400 μm × 400 μm white rectangle was used to quantify gene expression along the depth of the tree shrew SC, which was further divided into 14 subregions, each measuring 100 μm in length and 400 μm in width. In mice, the corresponding rectangle measured 800 μm × 250 μm and was divided into 16 subregions, each 50 μm in length and 250 μm in width.

To account for variations caused by animals and slices when comparing inhibitory and excitatory marker expression in tree shrews, we calculated the percentage of marker expression in each subregion relative to the total expression within the entire rectangle. This subregion percentage was then averaged across different slices from multiple animals. Next, we calculated the average total expression of inhibitory and excitatory markers across all slices within the entire rectangle. We then multiplied this average expression by the mean percentage for each subregion to obtain the mean number of inhibitory and excitatory marker expressions within each subregion. A similar analysis was conducted for mice, though vGAT and vGluT2 expression were detected in the same slices. Additionally, we confirmed that this analysis in mice produced the same distribution of inhibitory and excitatory neurons as the direct quantification of vGAT and vGluT2 in the same slices.

To assess the cellular co-localization of transcripts in tree shrews, we outlined 500 μm × 500 μm or 300 μm × 300 μm squares (Regions of Interest, ROIs) spaced from the middle to lateral regions at the surface or deeper layer of the SC. A similar approach was used in mice, where 300 μm × 300 μm squares were outlined from the middle to lateral regions for quantification.

## Notes

### Competing Interest Statement

The authors have declared no competing interest.

